# SARS-CoV-2 spike antibodies cross-react with dengue virus and enhance infection *in vitro* and *in vivo*

**DOI:** 10.1101/2023.10.09.557914

**Authors:** Kamini Jakhar, Sudipta Sonar, Gagandeep Singh, Tejeswara Rao Asuru, Garima Joshi, Nisha Beniwal, Tania Sarkar, Mahima Tiwari, Jaskaran Kaur, Deepak Kumar Rathore, Banwari Lal, Sandeep Kumar, Puneet Srivastav, Satendra Kumar, Vikas Phagna, Sushma Mithina, Lokesh Kumar, Vishal Gupta, Pallavi Kshetrapal, Savita Singh, Nitya Wadhwa, Ramachandran Thiruvengadam, Sreevatsan Raghavan, Mudita Gosain, Tripti Shrivastava, Sankar Bhattacharyya, Shailendra Asthana, Prasenjit Guchhait, Shailendra Mani

**Affiliations:** Centre for Virus Research, Therapeutics and Vaccines, Translational Health Science and Technology Institute, Faridabad, Haryana, India-120001; Computational Biophysics and CADD group, Computational and Mathematical Biology Centre (CMBC), Translational Health Science and Technology Institute (THSTI), Faridabad, Haryana, India-120001; Molecular Medicine, Regional Centre for Biotechnology, Faridabad, Haryana, India-120001; Centre for Immunobiology and Immunotherapy, Translational Health Science and Technology Institute, Faridabad, Haryana, India-120001; Translational Health Science and Technology Institute, Faridabad, Haryana, India-120001; Centre for Maternal and Child Health, Translational Health Science and Technology Institute, Faridabad, Haryana, India-120001

**Keywords:** SARS-CoV-2, Dengue Virus, Cross-reactivity, Antibody-Dependent Enhancement, AG129 Mice model

## Abstract

The presence of non-neutralizing antibodies of any dengue serotype, increase the severity of subsequent infection by other dengue serotypes. During SARS-CoV-2 pandemic, the number of symptomatic dengue cases increased in India. We describe that antibodies isolated from convalescent plasma from COVID-19 patients enhances DENV2 infection in vitro. CR3022, one antibody against SARS-CoV-2 spike protein, also showed elevated DENV2 infection in vitro. In silico protein-protein interactions between the spike antibodies and the DENV2 E-protein revealed significant interactions. Likewise, few monoclonal/polyclonal antibodies against SARS-CoV-2 showed increased dengue infection in vitro. Importantly, the AG129 mice infected with SARS-CoV-2 three-weeks prior to DENV2 infection, showed elevated dengue pathogenesis. Thus, highlighting the possibilities of elevated infection and symptomatic dengue disease in COVID-19 survivors.

## Introduction

Dengue is endemic to more than 120 countries with Asia contributing 70% of the global burden (1). Dengue virus (DENV) circulates as four serotypes (DENV 1-4) each containing multiple distinct genotypes. DENV infection can cause mild fever or lead to severe forms like Dengue Haemorrhagic Fever (DHF) and Dengue Shock Syndrome (DSS). Primary infection provides lifelong immunity to homotypic secondary infection but gives partial immunity from heterotypic challenge. Heterotypic secondary infections present with severe symptoms, due to higher viremia catalyzed by Antibody-dependent Enhancement (ADE). Sub-neutralizing antibodies that bind to heterotypic DENV promote FcR-mediated viral entry into cells (2). A stable attachment to Fc receptors by these antibodies, without reaching the neutralization threshold leads to ADE (3).

The dynamics of DENV infections during the SARS-CoV-2 pandemic vary globally. A 16% decline in dengue incidences was reported in 2020-21 compared to pre-COVID-19 (4). However, S. Khan *et. al.* revealed a substantial surge in dengue cases in 2021 in specific South Asian countries, including India (∼3-fold), Pakistan (>7-fold), and Bangladesh (>19-fold). This contrasted with a noteworthy decline of about 36% across Southeast Asia and Latin America (5). Several regional, environmental, and pandemic-related elements have been reported to shape these intricate dynamics. Disruptions in routine mosquito vector surveillance and control programs during COVID-19 lockdowns may have led to a rise in dengue cases, highlighting the crucial role of vector control in disease prevention. Similar pathology of DENV and COVID-19 presented diagnostic challenges, potentially contributing to underreporting (4) (6) (7) (8) (9).

Some studies indicate that ADE due to prior SARS-CoV-2 infection may elevate the risk of symptomatic and/or severe dengue. In Pakistan, a significant increase in dengue-related deaths (linked to DENV-2 prevalence) was observed in patients with a pre-exposure to SARS-CoV-2 (10).

Modest serological cross-reactivity between SARS-CoV-2 and DENV-2 has been reported (11) (12) (13). However, it is unclear if this would have any clinical impact on dengue cases (14). A study reported rabbit IgG against purified SARS-CoV-2 S1-RBD cross-reacts with DENV Envelope (E), precursor-membrane (PrM), and Non-structural protein 1 (NS1). However, these antibodies did not enhance DENV infection in THP-1 cells expressing FcR. Moreover, they impeded DENV infection at high concentrations (15). Another study showed *in vitro* inhibition of dengue infection by sera from convalescent COVID-19 patients’ sera (16).

Nath *et. al.* demonstrated serological cross-reactivity of SARS-CoV-2 positive serum samples with DENV1 and that the sera neutralized DENV1 (16). Anti-S1-RBD antibodies were shown to impede DENV infection at a specific concentration and COVID-19 patients’ sera exhibited a neutralizing ability against dengue infection (15). El-Qushayri *et al.,* in their systematic review, aimed to assess the clinical outcomes of patients with dengue and COVID-19 co-infection. They reported a high mortality rate in co-infected patients (19.1%), significantly exceeding the estimated global mortality rates for dengue (1.3%) and COVID-19 (2.04%). Also, they found that the hospital stay duration for co-infected individuals was longer than the reported median stays for standalone COVID-19 and dengue cases (17). Although, insightful, more information on the stoichiometry of the virus and antibodies in these studies, would have been useful, since it is crucial for understanding antibody-virus interactions and the potential for ADE Here, we evaluated the effect of preexisting SARS-CoV-2 antibodies on DENV-2 infection using cell-based assays and mouse model of DENV infection.

## Results

### *In silico* interaction mining drives the cross-recognition of DENV-E protein by anti-Spike and anti-RBD SARS-CoV-2 antibodies

To assess the cross-reactivity of SARS-CoV-2 Spike antibodies with DENV-2 E-protein dimer we used an *in silico* approach (Fig. 1A, B). We selected a panel of reported SARS-CoV-2 antibodies, based on their structure data (extent of their characterization, availability of the structure information of their co-crystals with low resolution, preferably below 4 A°.) and biological relevance (neutralization potential against SARS-CoV-2, target epitopes, etc.; Table-S1). For the identification of DENV-2 E epitopes, C8 antibody against E-protein dimer (Fab region, PDB-id: 4UTA) was used to characterize the interaction site based on interaction fingerprinting. For identifying the cross-reactive antibodies forming the most promising complexes with the E-protein; selected SARS-CoV-2 antibodies’ protein-protein (PP) blind docking was carried out with the whole DENV-2 E dimer. The consensus docked poses obtained from Piper and HDOCK tools were used for post-docking qualitative and quantitative analysis (pose selection and pose-filtering criteria). The top three SARS-CoV-2 antibodies forming the complexes with DENV-2 in terms of docking energy, interaction fingerprinting with co-crystal and MM-GBSA scores are CR3022 (Fab region, PDB-id:6W7Y), S2E12 ab (Fab region, PDB-id:7K45) and DMAb 2196 (Fab region, PDB:8D8R) (Fig. 1C).

**Figure 1.**
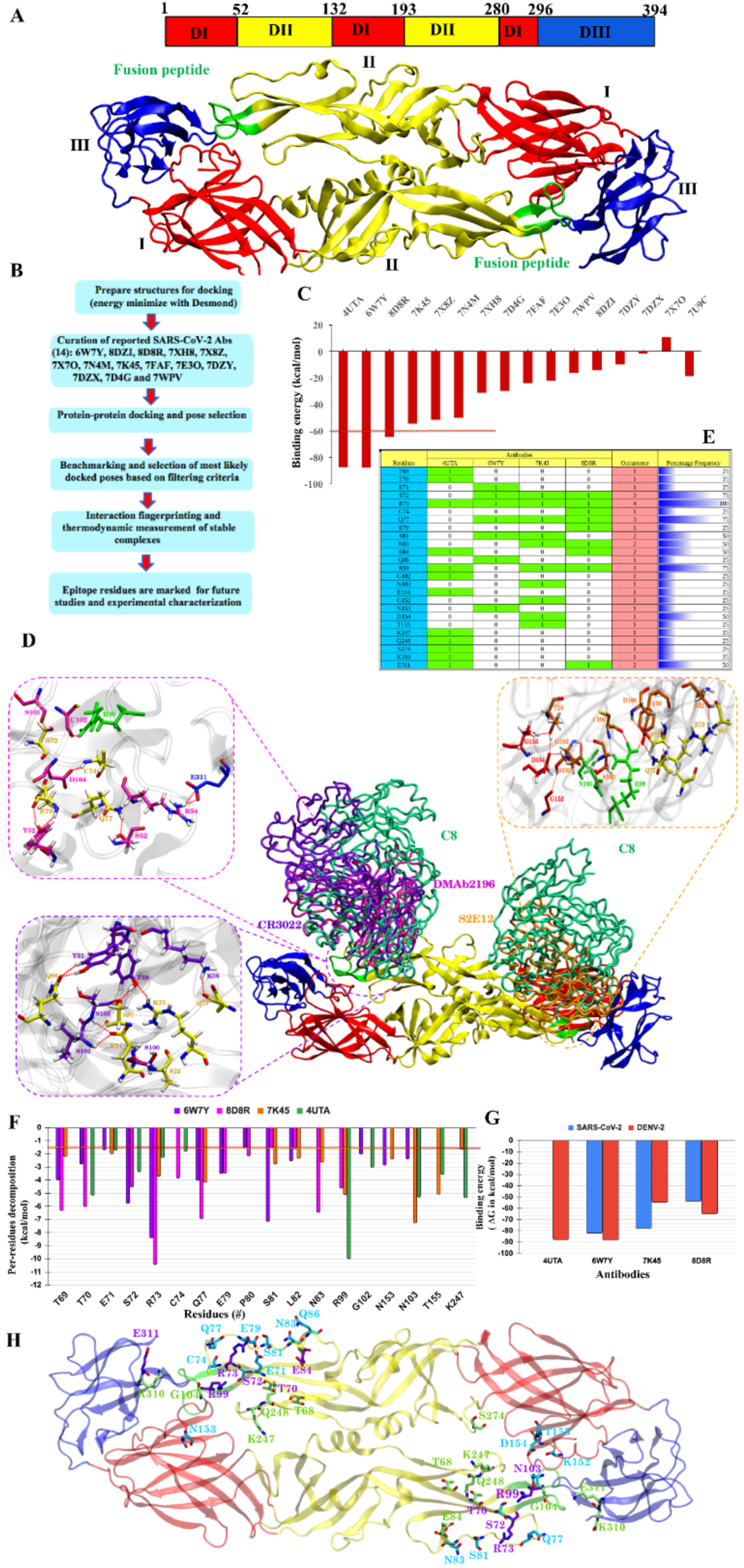
Computational insights exploring the cross-reactive SARS-CoV-2 antibodies at DENV-2 E protein: (A) Sequence and structural organization of the DENV-2 E dimer (PDB 1OAN). (B) Overview of the computational pipeline. (C) The binding free energy in kcal/mol of selected SARS-CoV-2 antibodies. The most likely complexes were generated using Protein-protein dockings. (D) The interaction map of top three antibodies (CR3022: 6W7Y: violet, S2E12 neutralizing antibody Fab fragment: 7K45: orange and DMAb 2196: 8D8R: magenta) along with the native antibody of DENV-2 E as control (C8:4UTA: green). The residue-wise zoom-out view is shown to highlight the critical residues mainly forming the interactions such as H bonds and SBs. The DENV-2 antibody interactions are shown in green-colored residues and SARS-CoV-2 antibody interactions are highlighted in the pink-colored residues. (E) The matrix of critical residues involved in either establishing either H-bonds or in SBs along with their occurrence. (F) The per-residue decomposition of top three antibodies indicating the thermodynamic contributions of epitope residues. (G) The net binding energy between identified antibodies and DENV-2 E which claims as potential in both cases. (H) The 3D positioning of identified epitope residues at dimer of DENV. The residues which only interact with SARS-CoV-2 antibodies are shown in cyan, the residues interacting with only DENV-2 antibody are shown in green color. The residues which are interacting with both DENV-2 E as well as with SARS-CoV-2 RBD are shown in violet color. All the residues are rendered in licorice and atom-wise C: cyan/green/violet, N: blue, O: red and S: yellow.

To quantitate their binding association at the residue level, major contributing amino acids were identified by residue-wise decomposition analysis setting the cut-off value of -1·5 kcal/mol. For the C8 antibody (4UTA) binding with DENV-2 E-protein, the interacting epitopes were majorly localized in Domain II (Fig. 1D). The binding sites of CR3022, S2E12 ab, and DMAb 2196 antibodies were found to overlap with C8 binding site (Fig. 1D). The contributing residues for all three antibodies were majorly located in Domain II including Fusion loop (Figure 1E, F). The findings of overlapping residues between C8 and the three SARS-CoV-2 interacting antibodies corroborate well with previously reported study (18). From the interaction analysis, the R73 appeared as the most common interacting residue in all antibodies. The pair-wise interaction analysis confirmed considerable overlapping residues between SARS-CoV-2 and DENV-2 antibodies (Figure 1E and 1F). Key residues that overlap between DENV-2 and at least two antibodies of SARS-CoV-2 are R73 (common among all), T70, S72 (Domain-II), R99 and N103 (Fusion loop) and E311 (Domain-III). We also performed the mining of paratope residues of SARS-CoV-2 antibodies with equal and/or increased interaction energy as C8 and identified overlapping residues between C8 and CR3022 (Figure S1).

Overall, the interaction maps and per-residue energetic decompositions showed that there were several common residues involved in the interaction of DENV-2-E with C8 and CR3022, S2E12 ab and DMAb 2196 antibodies indicating a high likelihood of cross-reactivity (Figure 1G and 1H, Table S2). The paratope residues involved in interaction of E-protein with CR3022 were also identified (Table-S3) and their interactions were assessed (Fig. S2).

### SARS-CoV-2 spike antibody CR3022 cross-reacts with DENV-2 and enhances DENV infection *in vitro*

Based on the computational outcomes, one of the antibodies, CR3022, which showed the highest affinity with DENV-2 E protein, was studied further. Based on the dengue virus susceptibility and cells monolayer robustness, we selected C6/36 cell line for confocal studies. Confocal microscopy images of CR3022 showed a statistically significant cross-reactivity with an average Mean Fluorescence Intensity (MFI) of 7·68 (p-value<0·0001) (Fig. 2A). Bio-Layer Interferometry (BLI), showed high affinity with a dissociation constant (KD) of <1·0pM with Spike protein (Fig. 2B(i)), and 2·83 µM with DENV-2 E-protein (Fig. 2B(ii)). Further, we observed a significant augmentation in the percentage of DENV-infected K562 cells with CR3022 antibody (p-value=0·01) in comparison to untreated virus control (Fig. 2C).

**Figure 2:**
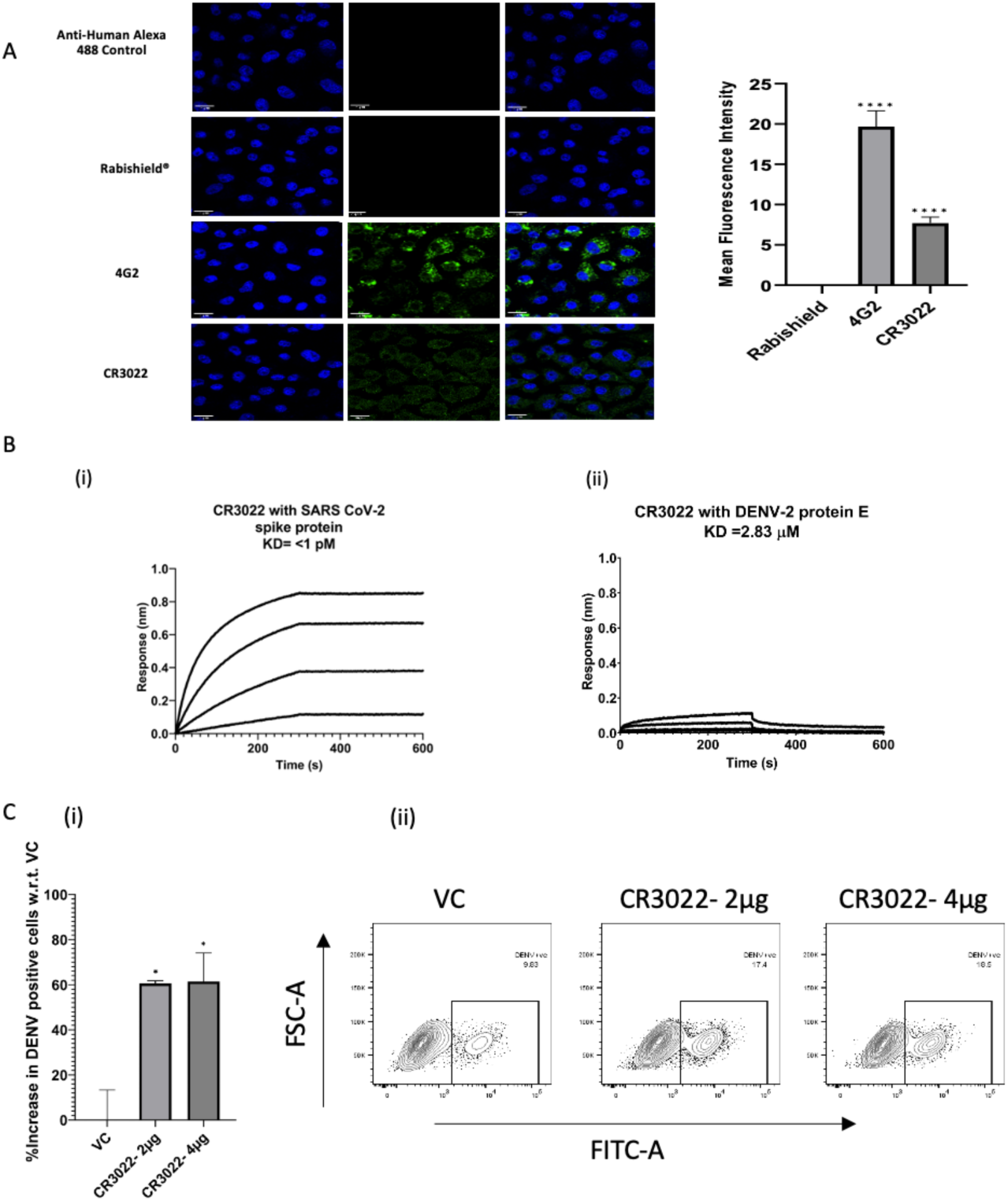
Cross-reactivity and ADE of Dengue virus infection by the SARS-CoV-2 anti-RBD antibody CR3022. (A) Confocal, digital images of DENV-2 (NGC) infected C6/36 cells stained with CR3022 along with the controls, 4G2 and Rabishield® (taken from Figure 1 to show the comparison) and the Mean Fluorescence Intensity (MFI). (B) Binding kinetics of (i) CR3022 with SARS-CoV-2 spike protein and (ii) DENV-2 protein E. (C) ADE of DENV-2 infection by different concentrations of CR3022 (2 µg and 4 µg) in K562 cells by flowcytometry. Statistical significance was checked using One-way ANOVA followed by Dunnett’s test. Asterisk (*) indicates a statistically significant difference between the control and treatment. P value = 0·1234 (ns), 0·0332 (*), 0·0021 (**), 0·0002 (***), <0·0001 (****).

### SARS-CoV-2 convalescent plasma enhances DENV infection *in vitro*

Samples from convalescent COVID-19 patients (n=48, Table-S4) from different time intervals of the pandemic were tested for their ability to cause ADE. We used FcγR-I, II and III expressing K562 and U937 cell lines. Both cell lines were shown to predominantly express FcγR-II (Fig. S3), as reported previously (19). The basal value of the only virus infection obtained in cell lines are normalized to the tested samples in order to calculate the fold change and graphs are plotted against virus infection (no serum) vs virus infection (incubated with serum) in tested samples. The DENV-2 infected K562 cells were observed to increase from 1 to 236 times in the first set of convalescent samples (recovered individuals during Wuhan wave), 77 to 235 times in the second set of convalescent samples (recovered individuals during/around the delta wave), and 44 to 172 times in the third set of convalescent samples (recovered individuals during omicron wave), compared to untreated virus control (Fig. 3A-C). The FACS gating strategy is given in Fig. S4. To confirm that the observed ADE effect was specifically mediated by antibodies present in the serum, IgG was purified from convalescent samples that exhibited high ADE activity. The assay was then performed using 10 µg of purified IgG and DENV-2 (clinical strain INDI-60). A significant enhancement of DENV-2 infection was observed in K562 cells incubated with purified IgGs from the convalescent samples #144 (P < 0.0001), #150 (P < 0.0001), and #151 (P < 0.0002), compared to the virus control, (Fig. 3D).

**Figure 3:**
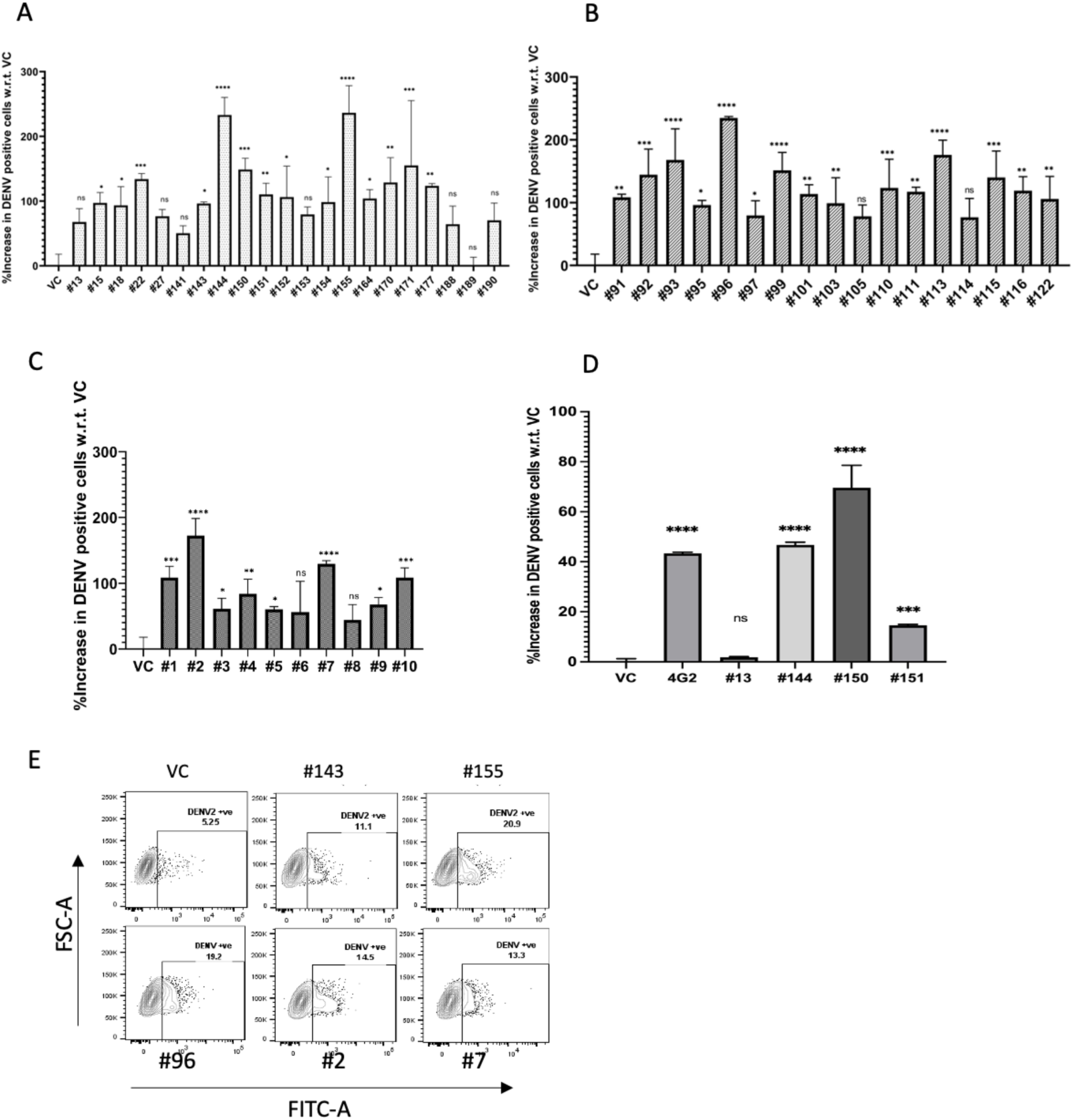
ADE of dengue virus infection by the SARS-CoV-2 positive patients’ sera in K562 cells. ADE due to convalescent plasma samples collected in the (A)first interval (Samples collected from May 2020-Jan 2021), n=21, (B) second interval (Samples collected from May 2021-June 2021), n=17, and (C) third interval (Samples collected from Feb 2022-April 2022), n=10. (D) IgG purified (10 µg) from the five convalescent samples showing the strongest ADE activity was assessed for its potential to enhance DENV infection using Dengue clinical strain (IND-60). The bar graph represents an average percentage increase in DENV positive cells w.r.t. VC for each sample with standard deviation for duplicates. Statistical significance was carried out using One-way ANOVA followed by Dunnett’s test in GraphPad Prism 8·4·2. Asterisk (*) indicates that the difference between the virus control and the serum samples is statistically significant. P value = 0·1234 (ns), 0·0332 (*), 0·0021 (**), 0·0002 (***), <0·0001 (****). (E) Representative dot plots of patient samples from the first interval (#143, #155), second interval (#96), and third interval (#2, #7). Data was analyzed on FlowJo version 10·8·1.

Using a subset of the convalescent plasma samples (n=15) similar results were observed in U937 cells, where DENV-2 infection was enhanced in the range of 32 to 292 times as compared to the virus control (Fig. S5A-B). A virus neutralization assay with SARS-CoV-2 Wuhan strain (Table-S5), did not show any association between the neutralizing antibody titer and enhancement in Dengue virus infection in convalescent plasma treated samples. (Fig. S6A-D). A representative quantitative PCR assay for intracellular DENV-2 genomic RNA showed a higher viral load in 70% of COVID-19 convalescent plasma treated samples as compared to the control (Fig. S7). This data correlates with the increase in DENV-2 positive cells observed in flow cytometry. Determination of the infectious titer of DENV produced from infected cells by focus-forming unit assay also showed a significant increase in the Foci-forming units/ml (FFU/ml) in COVID-19 convalescent plasma treated samples as compared to control (Fig. S8).

### SARS-CoV-2 antibodies enhance dengue virus infection *in vivo*

To validate the *in silico* and *in vitro* findings, we performed *in vivo* experiments where we took AG129 (Immunocompromised, lack of IFNAR1 and IFNGR1) mice within 6-8 weeks age group (N=21). The mice were divided into three groups, Group 1 as mock (G1, N=5), Group 2 as dengue virus control (G2, N=8), and Group 3 as test with both SARS-CoV-2 and dengue virus infection (G3, N=8). G2 was infected with dengue virus infection at day 21 and G3 was infected with SARS-CoV-2 virus at day 0 followed by dengue virus infection at day 21. Both G2 and G3 mice were infected with adeno-ACE-2 (Ad5CMVh, The University of Iowa Viral Vector Core Facility) viral vector at -5 days which is essential for SARS-CoV-2 infection. Body weight was observed for all the animals till day 27 when they were euthanized as shown in the schematic experimental plan (Fig. 4A). Average body weight in G2 mice group decreased at day 3 but it recovered with time (Fig. 4B). A significant decrease of body weight was observed in G3 as compared to G2.

**Figure 4:**
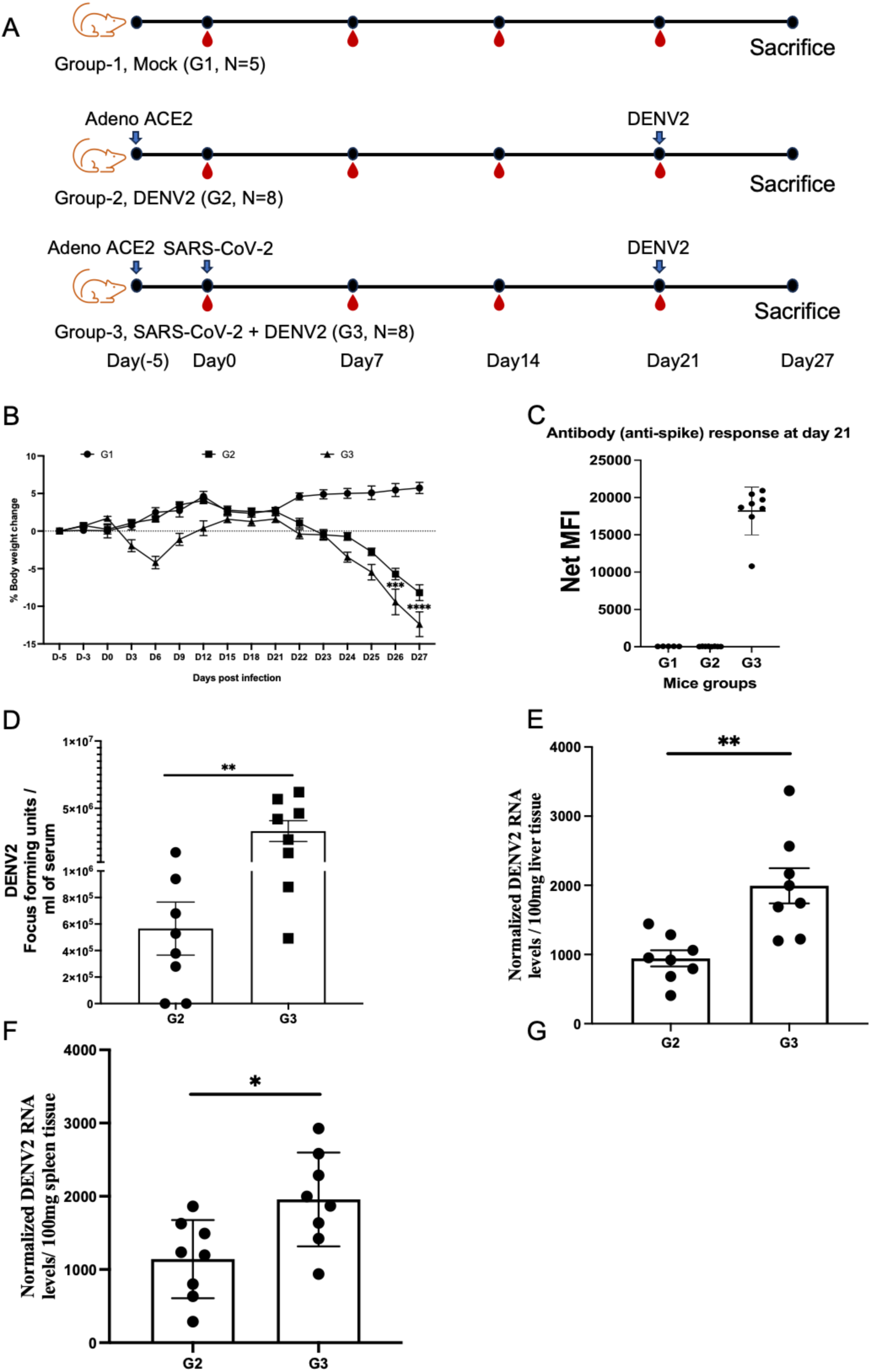
Effect of SARS-CoV-2 on dengue infection in AG129 mice: (A) Schematic experimental plan showing three animal groups: Mock as G1, adeno-ACE-2 viral vector infection at 5 days prior to day 0 in G2 and G3, SARS-CoV-2 infection at day 2 followed by infection with Dengue virus at day 21 in G2, and dengue virus infection at day 21 in G3. (B) Change in body weight. Two-way ANOVA was used. (C) SARS-CoV-2 Spike antibody response at day 21 (D) DENV2 viral particles were quantified by FFU assay in the serum of infected mice. To compare the two independent groups, the Mann-Whitney U test was applied for the statistical analysis. (E-F) The DENV2 genome was quantified by qRT-PCR from 100mg spleen and liver tissue. Based on the small sample size and comparing the two groups the Student T-test was applied and data are represented as mean ± SEM. *P < 0.05, **P < 0.01, ns = non-significant>< 0.05, **P < 0.01, ns = non-significant.

We further tested the antibody response against SARS-CoV-2 spike antigen at day 21 before DENV-2 infection, we observed the anti-spike IgG antibody response in all mice in G3 (Fig. 4C). Next, the live dengue virus particles determined in serum samples collected on day 27 (Fig. 4D), the data revealed a notable increase in number of viral particles in G3 mice group as compared to G2 mice group indicating enhancement of dengue infection with SARS-CoV-2 virus. Next, we tested the viral replication in the liver and spleen of the infected mice. We observed a high copy of viral RNA both in liver (Fig. 4E) and spleen (Fig. 4F) tissues. We also observed a decrease in the platelet count, as previously reported to occur during dengue infection. As shown in Fig (S9A), a decrease in the platelet count was seen in G1 vs G2 and G3 however, we did not observe any significant difference in G2 vs G3. WBC and Hematocrit measurement was also carried out but no significant differences in G2 vs G3 groups (Fig. S9A and S9B) were observed.

### SARS-CoV-2 spike antibodies cross-react with DENV-2

We wanted to check if the available sera/ antibodies cross react with DENV-2. To do that, SARS-CoV-2 anti-spike polyclonal antibody raised in rabbit (P1R) and mice (MSer) (obtained from BEI resources USA) were analyzed for cross-reactivity with DENV-2. Using confocal microscopy, we observed that both showed a significant cross-reactivity with an average (MFI) of 13·2 for P1R and 13·4 for MSer (Fig. 5A-B). Mouse Monoclonal antibodies against SARS-CoV-2, 1A9 (MFI of 7.4) and MM41 (MFI of 1.57), also showed cross-reactivity with DENV-2. The positive-control, 4G2 (pan-flavivirus-anti-envelope antibody), showed MFI of 18·28 whereas the Negative-control, anti-Rabies monoclonal (Rabishield®), showed no reactivity with DENV-2. The binding kinetics of 1A9 (SARS-CoV-2 spike antibody) showed a high affinity to DENV-2 Envelope with a dissociation constant (KD) of 4·84 µM (Fig. 5C ii-iii) using Bio-Layer Interferometry (BLI).

**Figure 5:**
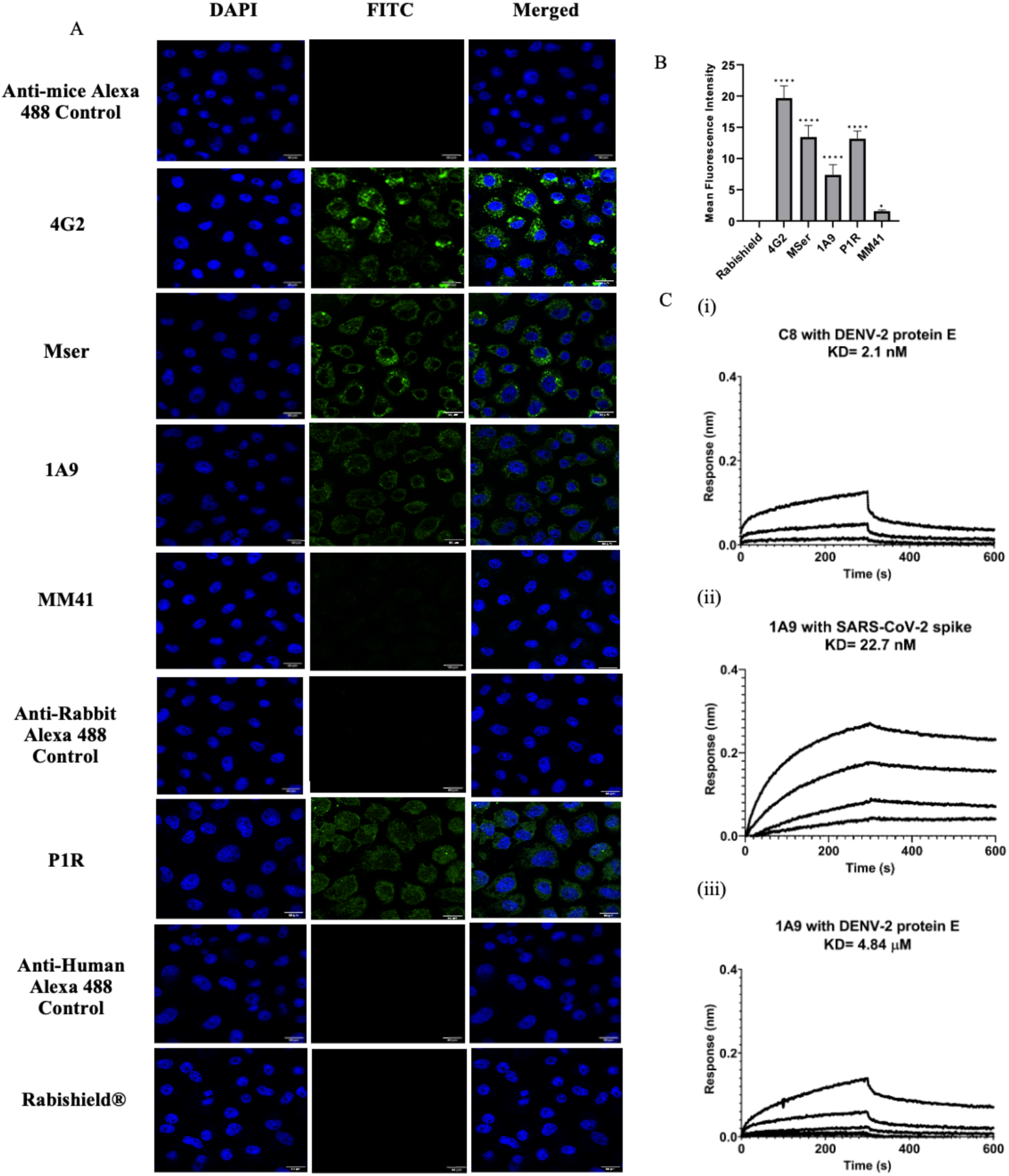
Cross-reactivity of antibodies and sera raised against SARS-CoV-2 Spike with DENV-2 (NGC) proteins. (A) Confocal, digital images of DENV-2 (NGC) infected C6/36 cells reacted with 4G2, antibodies and sera raised against SARS-CoV-2 after 48 hpi. Cell nuclei are stained with DAPI while the other antibodies are detected using FITC-labeled respective secondary antibodies. (B) Bar graph showing MFI of 4G2, MSer, 1A9, P1R (normalized w.r.t. the respective secondary antibody controls) with standard deviation. Analysis was performed using ImageJ and statistical significance was carried out using One-way ANOVA followed by Dunnett’s test in GraphPad Prism 8·4·2. Asterisk (*) indicates statistically significant difference in MFI w.r.t. Rabishield® (Negative control). P value = 0·1234 (ns), 0·0332 (*), 0·0021 (**), 0·0002 (***), <0·0001 (****). (C) Binding kinetics of C8 (DENV-E protein specific) and 1A9 (SARS-CoV-2 spike specific) antibodies with DENV-2 protein E and SARS-CoV-2 spike protein. Antibodies were loaded onto the sensors and serial dilutions of the proteins were used to study binding kinetics. The binding responses were calculated by subtracting data from reference and fitting globally with a 1:1 binding model using ForteBio’s Data Analysis software 10·1. The data was considered validated if 𝛘2 <0·5.

### SARS-CoV-2 spike antibodies enhance DENV-2 infection

Since DENV is endemic to India pre-existing anti-DENV antibodies may be responsible for the observed ADE. To rule out this possibility, we further tested commercially available monoclonal and polyclonal antibodies and sera raised in-house against the SARS-CoV-2 spike and RBD for their potential to enhance the DENV-2 infection (Table-S6). All of the tested antibodies (M1B, 1A9, MM41, P1R, M4B, M5B, Hser) enhanced DENV-2 infection (Fig. 6). The RBD specific antibody (M5B) showed stronger enhancement compared to the antibody against spike (MM41) with a dose-dependent increase in DENV-2 infection (Fig. 6A-B). Similarly, hamster serum against purified SARS-CoV-2 spike protein also showed ADE of DENV-2 infection in K562 cells (Fig. 6C). Monoclonal antibody against Rabies virus (Negative-control), did not show any ADE of dengue infection in K562 cells (Fig. 6C). A similar ADE of DENV infection was observed in U937 cells too (Fig. S10). Three monoclonal antibodies – 4G2, 1A9, and M4B, showed significant ADE, as determined by the infectious titer of the virus released in the culture supernatant from K562 cells (p-value<0·0001; Fig. S10). Having established that there is an enhancement of DENV-2 infection due to anti-SARS-CoV-2 antibodies, we next investigated whether it is time-dependent. We assessed ADE at 6 h, 18 h and 36 h post-infection (Fig.S11). Mice (Mser) and hamster (Hser) sera, showed significant ADE with respect to virus control (adjusted p-values Mser=0·0071 and Hser=0·0001), even at 6 hpi (Fig. S10). Although, we didn’t observe a significant enhancement due to 4G2 and anti-Spike monoclonals M3S and M2G at 6 hpi. At 18 hpi, a significant increase in the DENV-2 positive cells with respect to virus control was observed in 4G2 as well as M2G (Adjusted p-value=0·0101), which further enhanced significantly at 36 hpi (Adjusted p-values 4G2=0·0001 and M2G=0·0035). The ADE in Hser reached saturation as early as 6 h (Adjusted p-value<0·0001).

**Figure 6:**
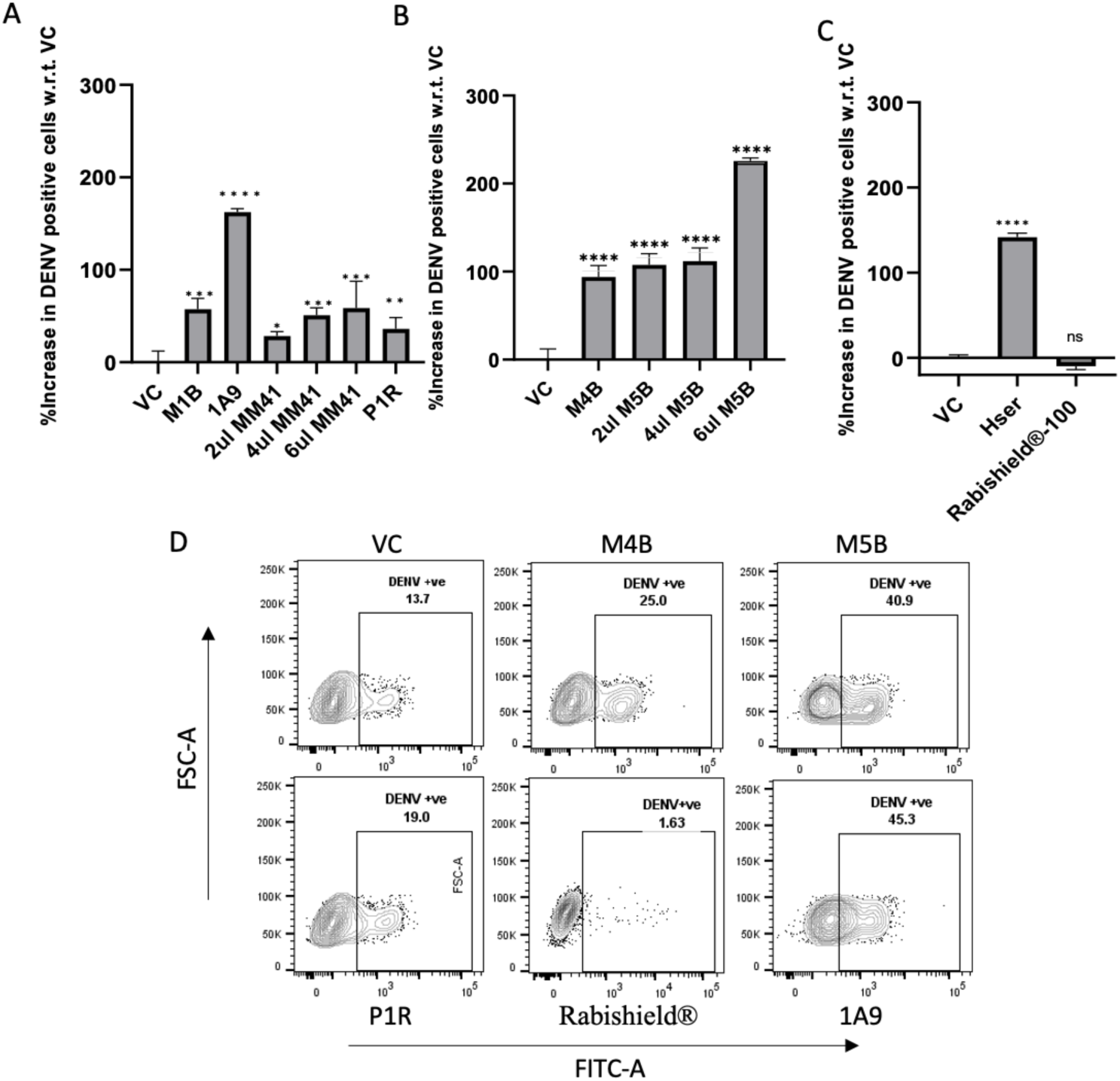
ADE of dengue virus infection by the monoclonal and polyclonal antibodies and serum against SARS-CoV-2. ADE due to (A) anti-spike monoclonal antibodies, M1B (5 µg), 1A9 (5 µg), and different volumes of MM41 (2 µg, 4 µg and 6 µg) and anti-spike polyclonal antibody raised in rabbit, P1R (5 µg), (B) anti-RBD monoclonal antibodies, M4B (5 µg), and different volumes of M5B (2 µg, 4 µg and 6 µg), (C) hamster serum immunized with SARS-CoV-2 spike and negative control (Rabishield®). Statistical significance was calculated using One-way ANOVA followed by Dunnett’s test in GraphPad Prism 8·4·2. Asterisk (*) indicates that the difference between the virus control and the serum samples is statistically significant. P value = 0·1234 (ns), 0·0332 (*), 0·0021 (**), 0·0002 (***), <0·0001 (****). (D) Representative dot plots. Data was analyzed using FlowJo version 10·8·1.

## Discussion

In this study, we demonstrate that SARS-CoV-2 antibodies cross-react with DENV-2 and enhance DENV infection. A thorough *in silico* study indicated the high likeliness of cross-reactivity of SARS-CoV-2 spike antibodies with DENV-2 E protein. SARS-CoV-2 spike protein is critical for viral entry into cells and is the major target not only for neutralizing antibodies but also for infection-enhancing antibodies (20). The *in silico* results indicated that there is a significant interaction of SARS-CoV-2 antibodies with DENV-2 dimeric E protein. The binding free energy of SARS-CoV-2 antibodies for example, CR3022 (6W7Y), S2E12 ab (7K45) and DMAb2196 (8D8R), being comparable to the DENV-2 dimer specific antibody, C8 (4UTA). These top three binders (SARS-CoV-2 specific antibodies) shared their interacting interface with C8 antibody. The residues involved in the interaction of C8 antibody with DENV-2 E, were also involved in the interaction of at-least one of the top three binders with E-protein. The contributing residues which were found to be majorly stabilizing the interaction of these top three SARS-CoV-2 antibodies with E-protein. We also observed that convalescent plasma samples of SARS-CoV-2 recovered individuals, collected at three different time intervals during COVID 19 pandemic, significantly enhanced the DENV-2 infection in cultured Fc-receptor positive K562 and U937 cells. We observed an increased viral load as indicated by FFU and also increased viral RNA levels in spleen and liver in AG129 mice group which had SARS-CoV-2 antibodies. The finding in animals model strongly supported our *in silico* and *in vitro* data which clearly showed that SARS-CoV-2 spike antibodies mediated enhancement of dengue virus infection.

Our work supports the work published by others highlighting SARS-CoV-2 cross-reactivity with dengue virus and false positivity with dengue serology (21) (22). We used the DENV-2 envelope protein for BLI and computational studies since it has been reported to have maximum cross-reactivity with SARS-CoV-2 S1 specific antibodies (11). The dimeric DENV-2 E-protein was used for docking studies as we intended to study the interaction of DENV-2 with SARS-CoV-2 spike antibodies, which would happen before the viral entry, extracellularly and it’s known that the E-protein exists as a head-to-tail oriented homodimer on the surface of mature-virions (23). All the residues involved in the interaction of the top three binders of SARS-CoV-2 antibodies with E-protein when mapped, were majorly found to lie in the Domain-II and fusion-loop of E-protein. It is well known that the antibodies against two epitopes of DENV are strong mediators of ADE: prM epitopes (24) and fusion loop epitopes (25). The fusion loop epitopes could be the ones, potentially involved in ADE. Multiple residues have been identified from *in silico* interaction mining, among them few residues such as R73 (common among all), T70, S72 (Domain-II), R99 and N103 (Fusion loop) and E311 (Domain-III) seem critical. These residues put forth the possible avenue to explore further for epitope-based therapeutic interventions. The BLI findings between dengue envelope protein and antibodies against SARS-CoV-2 spike protein indicate a strong interaction with KD values in µM range.

The DENV-2 used for the *in vitro* assay was prepared in C6/36 cells and wasn’t characterized for the proportions of immature and mature-virions. It is well known that the virus prepared *in vitro* in mosquito cells has a considerably greater number of immature virions as compared to mammalian cells, probably due to differential furin-expression (26) (27). Also, it is recognized that laboratory adapted DENV strains are more dynamic than clinical strains and expose cryptic epitopes more easily (28). Thus our virus preparation might have major proportion of immature virions but this probably won’t change the fact that the ADE of DENV-infection would happen since the viral heterogeneity in response to the natural DENV-infections is also well known The antibody repertoire in the natural infections is known to have an abundance of poorly neutralizing antibodies which are majorly generated in response to the immature conformations (29), which is a result of incomplete furin processing, resulting in egress of immature-forms along with and mature-forms of virions post-assembly (30) (31). Also, the fully mature-virions have been reported to be less susceptible to neutralization by cross-reactive and heterotypic immune sera than partially mature variants (29). Thus, the heterogeneity in the DENV-2 preparation used for the *in vitro* assays can be probably closely related to the natural infection scenario. Although we did not have the history of patient’s (SARS-CoV-2 Convalescent samples) earlier infection from other dengue serotype which could have resulted in ADE in our finding, we tested the commercially available SARS-CoV-2 antibodies and Spike-immunized mice and hamster sera. Monoclonal and polyclonal antibodies raised against SARS-CoV-2 showed an increased DENV-2 infection in both cell-based models which re-confirms our observations. Due to limitation on dengue virus serotypes, we only tested ADE against Dengue serotype 2 (NGC strain and Clinical isolate (Indi-60) strain) but we believe that we will have similar ADE effect as all the dengue serotypes are very closely antigenically related. However, in the animal experiment, the platelet, WBC counts and hematocrit levels were not significantly affected between mice given dengue (G2) and SARS CoV-2 + DENV (G3). As these parameters are associated with disease severity which could explain why we did not observe dengue severity during the SARS-CoV-2 pandemic which are promoting us to dig more deeper and further studies to find the missing links. We also observed an increase in dengue viral load in G3 (SARS CoV-2 + DENV infected) group mice as compared to G2 (dengue infected) group mice, that might be indicating an increase in the number of the symptomatic cases in endemic population rather asymptomatic cases. It is still not clear if there was underreporting or reduced testing for dengue during the peak SARS-CoV-2 pandemic as the clinical symptoms of both viruses are very similar. Our understanding is still limited and future studies might shed some light on the this.

The limitation of the study is the availability of the pre-pandemic clinical samples like plasma/Serum or PBMCs, which could have provided a better insight and might have shed some light on why these all are not leading to dengue severity in the real-world scenario. We tried to overcome this using the animal model but a clinical sample would have been better comparator.

Based on the current finding from *in silico, in vitro and in vivo* experiments, the study suggest that SARS-CoV-2 spike directed antibodies cross reacts with dengue envelope protein and enhance dengue viral load in AG129 experimental mice. Our findings signify the importance of the cross reactivity and viral interactions in endemic regions where more than one virus is circulating at the same time. Understanding the viral interactions during the co-circulation/Co-infections needed more research in future to understand how they pose vaccine development challenges and will impact on public health. These things will also pose significant challenges on surveillance and/or vaccine running programs which might provide a false positivity due to viral cross reactivity. So having an assay which can clearly differentiate the viral infection will be more beneficial approach in future.

## Materials and Methods

### Molecular docking of DENV-2 Envelope glycoprotein and SARS-CoV-2 antibodies

#### Curation of data

X-ray crystal structures of DENV-2 envelope glycoprotein (PDB ID:4UT6) (32) and SARS-CoV-2 antibodies were curated from RCSB in PDB format (http://www.rcsb.org/pdb). These crystal structures along with their corresponding resolution values, and are summarized in – Table-S1. Crystal structure of antibodies were curated by removal of spike protein, water molecules and metal ions using UCSF Chimera 1·15 (33). All structures were prepared using Maestro’s Protein Preparation Wizard module (Schrödinger Release 2020-1) (34). The hydrogens and bond orders were added, missing side-chains and loops were filled by Prime (34). The hydrogen bond (HB) optimization and restrained minimizations were also done for the systems using force field OPLS4 (35).

#### Protein-Antibody docking and choosing the best pose

Protein-protein interactions govern various aspects of structural and functional cellular mechanisms, and their elucidation is crucial for a better understanding of biological processes. All the prepared SARS-CoV-2 antibodies, along with controls i.e positive (pdb-id:4UTA) and negative control (pdb-id:7U9G) were docked on prepared DENV2 E protein structure by masking non-CDR regions of antibodies. The Chothia definition is used for the CDR region. Multiple docking methods were used to minimize the biases by including HDOCK (36), pyDock (37) and Piper program (38). The rationale for implementing multiple docking programs was to obtain the consensus results from all three tools and to remove any false-positive outcomes. Best poses were chosen from docked models on the basis of their docking score, cluster size (39) and reported paratope residues (40) (41) (42).

### Thermodynamic quantification using Molecular Mechanics-Generalized Born Surface Area (MM-GBSA)

The binding free energy of protein and antibody complexes was carried out in Prime MM-GBSA (Molecular Mechanics Generalized Born Surface Area) by Prime (43) module of Schrodinger suite. We used the VSGB solvation model and OPLS4 force field. The binding free energy is calculated by: MMGBSA dG Bind (NS) = Complex – Receptor (from optimized complex) – Ligand (from optimized complex) = MMGBSA dG Bind–Rec Strain–Lig Strain, Where, Rec Strain = Receptor (from optimized complex) – Receptor, Lig Strain = Ligand (from optimized complex) – Ligand, MMGBSA dG Bind = Complex – Receptor – Ligand, NS = no strain.

### Figures

All figures were generated using VMD (V.1·9·1) (44) and graphs were generated using the XMGRACE (Version 5·1·19) (42).

### Cells

Human myeloid cell lines K562 (ATCC CCL-243) and U937 (ATCC CRL-1593.2) were cultured in Iscove’s Modified Dulbecco’s media (IMDM, HIMEDIA) and Roswell Park Memorial Institute (RPMI, Gibco) 1640 respectively. African Green Monkey cell line Vero E6 (NCCS, India) was maintained in Dulbecco’s Modified Eagle Medium (DMEM, HIMEDIA) while A. albopictus cell line C6/36 (NCCS, India) was cultivated in Leibovitz’s L-15 Medium (HIMEDIA). All the cells were maintained in respective media supplemented with 100 U/ml of penicillin, 100 μg/ml of streptomycin (HIMEDIA), and 10% Fetal Bovine Serum (HIMEDIA). C6/36 cells were maintained at 28°C (ESCO incubator) without CO_2_ while the other cells were grown in a humidified 37°C incubator (Thermo Fisher Scientific) with 5% CO_2_.

### Virus stocks

Dengue virus serotype 2 (NGC strain) and (P8-P23085 INDI-60) were propagated in C6/36 cells and it was used for all the assays. The cells at a confluency of 60-70% were infected with Dengue virus at a multiplicity of infection (MOI) of 0·1 using inoculum prepared in L-15 medium supplemented with 2% FBS. It was incubated with cells for 2 h at 28°C with intermittent rocking. Subsequently, the cells were washed and incubated with L-15 medium with 10% FBS. Culture supernatant was harvested at 5 dpi, clarified by centrifugation at 129 ×*g* at room temperature and concentrated using centrifugal ultrafiltration (Amicon® Ultra centrifugal filter (100 kDa), ACS510024 from Millipore, Darmstadt, Germany) at 3220 ×*g* for 10 min for every 15 ml of supernatant. The infectious titer of the virus was determined by a focus-forming unit (FFU) assay performed in Vero E6 cells.

The three variants of SARS-CoV-2, original Wuhan (USA-WA1/2020, Genbank: MN985325), B·1·617·2 (δ-variant) known as THSTI_287 (GenBank: MZ356566·1), and Omicron BA·1(GISAID accession no.: EPI_ISL_8764350) were propagated in Vero E6 cells. Cells were infected with 0·1 MOI of virus inoculum and incubated for 1 h with shaking at 37°C. Virus supernatant was harvested at 48 hpi, filtered, and stored at -80°C. The infective titer was determined by titration on Vero E6 cells using the TCID_50_ method (45). These virus stocks were used for neutralization assays.

### Serum samples and Antibodies

We selected a group of convalescent plasma samples (n=48) collected at three different time intervals during COVID-19 pandemic in India. They are defined as follows: the first interval was from May 2020 to January 2021, second interval from May 2021 to June 2021, third interval from February 2022 to April 2022. The clinical details of the samples are given in the Table S1. The pan flavivirus, anti-envelope mAb, 4G2 was purified inhouse from HB112 hybridoma cells. SARS-CoV-2 spike (S2 specific) antibody 1A9 was purchased from Gentex while Anti-Recombinant SARS-CoV-2 Spike Protein (whole spike) Sino 4059-MM41 was purchased from Sino Biologicals. Other SARS CoV2-related antibodies M1B, M4B, M5B, P1R were obtained from BEI resources, USA. SARS CoV2 infected mice and hamster serum were generated inhouse. Rabies Human monoclonal antibody, Rabishield®-100 (Serum Institute of India), was purchased from the pharmacy. All the details regarding commercial antibodies/ serum are mentioned in-Table S2.

### Confocal microscopy

0.5 x 10^6^ C6/36 cells from an exponentially growing culture were seeded on grass cover slips followed by DENV-2 at 1.0 MOI. At 48 h post-infection the cells were fixed with 3.7% paraformaldehyde (PFA) and permeabilized using 0.1% Triton-X-100. The cells were immune-stained with primary antibody and fluor-conjugated secondary antibodies. Subsequently, coverslips were mounted onto glass slides using ProLong Gold Antifade Mountant supplemented with DAPI. Fluorescence was observed under 60X magnification using a FLUOVIEW FV3000 confocal microscope (Olympus). The mean fluorescence intensity (MFI) was calculated using images from 10 fields per slide with 7 cells per field (at least 70 data points). Statistical analysis employed One-way ANOVA followed by Dunnett’s test, comparing to the negative control (Rabishield).

### Binding kinetics using Biolayer Interferometry (BLI)

BLI studies were performed on an Octet RED 96 instrument at 24°C with shaking at 1000 rpm (ForteBIO). For DENV-2 Envelope protein, the buffer composition was 50mM Tris, 150mM NaCl, 0.5% glycerol, pH 8.0. Whereas, for SARS-CoV-2 Spike protein, phosphate buffered saline, pH 7.4 was used. Sensors were in the respective kinetic buffer (46). For studying binding kinetics with protein E, antibodies at a concentration of 5 µg/ml were loaded onto the sensors (AHC (Anti-hIgG Fc Capture) for CR3022 and C8, protein G (for 1A9 antibody)) DENV E dimeric protein was diluted serially three-fold (starting with 6000nM) and Spike trimeric protein was diluted 2-fold (starting with 600nM) in their respective kinetic buffers. The designed experiment started with a baseline (100Sec), followed by 3 cycles of neutralization and regeneration and again a baseline (100Sec). After this step, antibody loading was performed (700sec) onto all the 5 sensors. After the loading step, another baseline (baseline2, 60sec), Association was performed after the 2nd baseline step for 300sec, followed by 300sec dissociation.

Here the 5th sensor was used as reference where no analyte was used, where the analyte concentration was kept 0nM. The same sensor during the data processing was assigned as reference well (change well type-reference well) and same wells; reference well, were subtracted from the raw data. Here the ligand and analyte dilution as well as the whole experiment was performed in the respective kinetic buffers. The binding responses were calculated and fitting globally with a 1:1 binding model using ForteBio’s Data Analysis software 10·1. The data was considered validated if X^2^ <0·5.

### Antibody-dependent Enhancement (ADE) Assay

Convalescent Serum samples (15 μl), Purified IgG antibodies (10 µg) and commercial antibodies (at various concentrations) were diluted in plain Iscove’s Modified Dulbecco’s Medium (IMDM) to a total volume of 100 μl. Subsequently, 100 μl of virus dilution was added, and the mixture was incubated at 37°C for 1 h. The virus-antibody complexes were then allowed to infect 1 x 10^6^ cells K562/U937 cells for 2 h at an MOI 1. After infection the cells were washed and incubated in IMDM-2% FBS at 37°C, 5% CO_2_ for 36 h. Subsequently, cells were fixed using 4% PFA, permeabilized with 0.1% Triton X-100, immune-stained with purified 4G2 (dilution 1:200), and Alexa-Fluor 488 conjugated anti-mouse (dilution 1:500). The number of cells positive for DENV antigen was quantified by flow cytometry.

### Isolation and Purification of IgG from Serum samples

Serum sample was centrifuged at 10,000 x g at 4 °C for 10 min followed by adding 1 part of 1 M Tris-HCl, pH 8.0 to 10 parts of serum sample, to adjust the pH. Drop by drop, saturated ammonium sulphate solution up to 45% saturation was added to it and it was stirred for 1 h. Serum sample was again centrifuged at 10,000 x g at 4 °C for 10 min and supernatant was discarded. Pellet was dissolved in minimum volume of the desalting column buffer. Sephadex G-25 column was once cleaned with distilled water and then equilibrated with 10-15 ml of binding buffer. The sample was loaded onto the column and flowthrough was collected in a fresh 15ml falcon. Binding buffer (5-10 ml) was again passed through the column and about 1-2 ml of flowthrough was collected in the same falcon tube. To separate the IgG, protein G column was rinsed with ethanol and then distilled water. Collection tubes with 50 μl of 1.0 M dibasic sodium phosphate was prepared to ensure that the final pH of the collected samples is close to neutral. The packed column was washed with five column volumes (∼25 ml) distilled water to remove the slurry storage medium. Column was equilibrated with five to ten column volumes of 50mM sodium phosphate, pH 7.0 and 500 mM NaCl (binding buffer). Pre-treated sample was loaded on the equilibrated column. Column was washed with∼ 25 ml binding buffer followed by elution with 5 column volumes of elution buffer. Fractions of 500 μl were collected into the collection tubes containing the neutralizing buffer. The fractions were analysed by absorbance at 280 nm in a spectrophotometer to measure protein concentration and SDS-PAGE to confirm purity of IgG.

### Flow Cytometry Analysis

Flow cytometry was conducted using a BD FACS Canto II flow cytometer (BD Biosciences) with 20,000 cells acquired per sample through FACS Diva software. FlowJo v10.8.1 was used for analysis. Live cells were gated based on FSC and SSC, followed by singlet determination using FSC-A vs FSC-H gate. Dengue virus-positive cells were gated on the FITC channel using singlet population. FITC voltage was set using isotype control. Fcγ receptor expression was assessed with anti-CD64 (Alexa Fluor 700), anti-CD32 (PE), and anti-CD16 (APC) antibodies in K562 and U937 cells. Statistical analysis was conducted in GraphPad Prism 8.

### Live Virus Neutralization Assay

Two-fold serial dilutions of heat-inactivated plasma samples were mixed with 100 TCID_50_ of SARS-CoV-2 virus and subsequently allowed to infect Vero E6 monolayers. After incubation for 72 h at 37°C with 5% CO_2_, the degree of CPE was qualitatively determined under a microscope. Absence of CPE indicated virus neutralization, and the highest dilution that completely prevented emergence of CPE was considered as the neutralization titer. For plasma samples collected during the third interval, cells were fixed 24 h post-infection and immune-stained using antibody against the SARS-CoV-2 Nucleocapsid protein (1:2000, 4H2, Genscript), followed by Alexa 488 conjugated anti-mouse antibody (1:500, Invitrogen). Fluorescent foci were quantified by AID iSpot Reader with AID EliSpot 8.0 iSpot software.

### Detection of DENV RNA using Quantitative RT-PCR

Total RNA from 1 x 10^6^ K562 cells was extracted using the standard TRIzol method. RNA was quantitated using Nanodrop (Thermofisher Scientific) and 300 ng RNA was used for qPCR using a one-step RT-PCR kit (SOLIScript® 1-step Probe kit, Solis Biodyne) following the manufacturer’s protocol for detection of DENV2 RNA. GAPDH was used as an internal control. Fold change in DENV2 RNA copies was calculated with respect to GAPDH. The details regarding primer and probe sequences are given in Table S7.

### Focus forming unit (FFU) assay

FFU assay was carried out to determine the virus titer in the serum of DENV2 infected AG129 mice with slight modifications. Vero E6 cells were seeded at a density of 0.01x10^6^ cells/well in a 96-well black/clear bottom plate and cells were allowed to reach 80% confluency. At 80% confluency 100 µl of 10-fold serially diluted serum in media with 2% FBS was added to the wells and incubated. After 2 hr, serum was removed, fresh media with 2% FBS was added and incubated for 24 hr at 37^°^ C in 5% CO_2_. Following that cells were fixed with 2% PFA and the immunostaining for virus detection was performed using mAb 4G2 (pan-DENV anti-envelope antibody), followed by secondary antibody, goat anti-mouse IgG, AF488 conjugated (Invitrogen) prepared in permeabilization buffer (0.1% triton x100 with 2%BSA in PBS). The foci were visualized and counted using fluorescence microscope.

### Mouse Infection

The AG129 (IFNα/β/γR -/-129/Sv) mice were obtained from the Jackson laboratory, USA and maintained in small animal facility (SAF), RCB, Faridabad, India. All experiments with AG129 mice were performed in 6-8 weeks old mice of either sex. One group of mice were treated (one-time through intranasal route) with human ACE2 adenovirus using 2.5x10^8^ PFU for transient expression of hACE2 in lungs and respiratory track. After five days mice were infected with SARS-CoV-2 via nasal route inoculation using 1x10^5^ PFU virus. Mice were bled at day7, 14 and 21 post SARS-CoV-2 infection to measure the antibodies against SARS-CoV-2. At day 21 both the groups were challenged with 1x10^4^ FFU mouse-adapted DENV2 (P8-P23085 INDI-60) and were euthanized on the moribund state.

### Quantitative real time PCR

Total RNA from the mice tissues was extracted using RNAiso (Takara Bio, Japan) followed by phenol-chloroform treatment. The first strand cDNA was synthesized from 1µg RNA using cDNA synthesis kit (BioRad, USA) as per manufacturer’s protocol. The cDNA was used for real-time PCR using SYBR green supermix (BioRad) in an Applied Biosystems Quant Studio^TM^ 6 Flex Real-Time PCR System. The primers set used for detection of the respective genes are given in Table S8.

### Statistical analysis

Experimental values were represented as mean ± standard deviation (SD). Statistical analysis was done using the ordinary one-way ANOVA followed by Dunnett’s multiple comparison test, using GraphPad Prism version 8·4·2. Differences were considered significant at p-value < 0·05.

## Acknowledgements

The following reagent was obtained through BEI Resources, NIAID, NIH: SARS-Related Coronavirus 2, Isolate USA-WA1/2020, NR-52281, Monoclonal Anti-SARS-Related Coronavirus 2 Spike Glycoprotein, Clone 1-3D7 (produced *in vitro*), NR-56488, Monoclonal Anti-SARS-Related Coronavirus 2 Spike Glycoprotein Receptor Binding Domain-(RBD), Clone 2TP1C3 (produced *in vitro*), NR-55296, Polyclonal Anti-SARS-Related Coronavirus 2 Spike Glycoprotein (IgG, Rabbit), NR-52947.” The RBD is proprietary reagents with IP No. 202011018845, we thank Dr. Barney Graham (VRC/NIAID/NIH) for providing us with spike construct (SARS-2-CoV S 2P). We would like to thank Prof. Shinjini Bhatnagar, Prof. Pramod Garg and Prof. Jayanta Bhattacharya from THSTI for critically reviewing the manuscript, ABSL3 facility committee and staff for their help and support. This research has been conducted with the significant contribution and expertise of THSTI, NCR Biotech Science Cluster BIOREPOSITORY and DBT India Consortium”.

DBT India Consortium**, ** Name of all the collaborating Institutes/ Hospitals: Translational Health Science Technology Institute, Clinical partners: Maulana Azad Medical College, Lok Nayak Jai Prakash Hospital, and Lady Hardinge Medical College in Delhi, ESI Medical College Hospital, Faridabad, Civil Hospital, Gurugram, Haryana; Civil Hospital, Palwal, Haryana; Al-Falah School of Medical Science & Research Centre and Hospital, Dhauj, Haryana; Medanta Hospital, Gurugram; Shaheed Hasan Khan Mewati Government Medical College, Nalhar, Haryana.

## Data availability

All the data are presented in this manuscript. The virus strains used in this study have the following accession numbers: original Wuhan (USA-WA1/2020, Genbank: MN985325), B·1·617·2 (δ-variant) known as THSTI_287 (GenBank: MZ356566·1), and Omicron BA·1 (GISAID accession no.: EPI_ISL_8764350) Source data are provided with this paper.

## Author Contributions

SM conceptualized the idea. SM, SB, SA, TrS and PG coordinated the research activity. KJ and SS contributed to designing and execution of viral assays. KJ, SS, GS, NB, TS, MT, JK, DKR, BL, SK, PS, SK, VP, S, LK and VG contributed to data curation, investigation and methodology. KJ, SS, TRA, GJ, GS, SB, SA, TrS, PG and SM contributed to formal analysis. SS, SR, MG, PK, NW, RT, SB, TrS, SA and SM provided resources. KJ, SS, TRA, GJ, TrS, PK, NW, RT, SB, SA, PG and SM contributed to writing, review & editing the draft. All authors had full access to all the data in the study and had final responsibility for the decision to submit for publication.

## Ethics statement

We obtained the approval from the Institutional Bio-Safety Committee and Review Committee on Genetic Manipulation (RCGM), for *in vitro* and *in vivo* work, prior to conducting the studies: (IBSC#495/2023), RCGM (No.-BT/IBKP/137/2020), RCGM (No.BT/IBKP/053/ 2020).

