## Supplementary file for "SARS-CoV-2 spike antibodies cross-react with dengue virus and enhance infection *in vitro* and *in vivo*"

### Supplementary Data

**Appendix**

**Supplementary Table S1: Details of antibodies for computational study -------------------04**

**Supplementary Table S2: Details of common epitopes indicated by computational study08**

**Supplementary Table S3: Details of paratope residues involved in interaction of E-protein with CR3022-----------------------------------------------------------------------------------------09**

**Supplementary Table S4: Clinical data of the 48 patient samples taken for study--------10**

**Supplementary Table S5: Neutralizing antibody titers of the commercial SARS-CoV-2 antibodies, animal sera, and convalescent plasma samples against the original Wuhan strain---------------------------------------------------------------------------11**

**Supplementary Table S6: Details of antibodies and sera used for the study---------------15**

**Supplementary Table S7: Primer and probe sequences used of DENV and GAPDH in qRT PCR-----------------------------------------------------------------------------------------17**

**Supplementary Table S8: Primer and probe sequences used in qRT PCR for animal tissues-----------------------------------------------------------------------------------------------18**

**Supplementary Figure S1: Mining of antibodies interacting residues-----------------------19**

**Supplementary Figure S2: Protein interaction diagram showing paratope residues of mAb CR3022 (chain H L) involved in interaction with E-protein of SARS-CoV-2 (Chain A B).------------------------------------------------------------------------------------------------20**

**Supplementary Figure S3: Expression of Fcγ Receptors (FcγRs), type I, II, and III in undifferentiated K562 and U937 cells--------------------------------------------------------------21**

**Supplementary Figure S4: Gating strategy for detection of DENV-positive cells in flowcytometry---------------------------------------------------------------------------------------22**

**Supplementary Figure S5: ADE of dengue virus infection by the SARS-CoV-2 positive patients' samples in U937 cells-----------------------------------------------------------23**

**Supplementary Figure S6: Comparison of neutralizing antibody titres of patient samples against SARS-CoV-2 Wuhan with the increase in dengue virus infection--------24**

**Supplementary Figure S7: Quantitation of dengue virus ADE using qRT PCR-----------25**

**Supplementary Figure S8: Replicating virus in supernatants from ADE of dengue by the antibodies and convalescent plasma samples against SARS-CoV-2 in K562 cells---------26**

**Supplementary Figure S9. Haematological analysis of dengue-infected AG129 mice following SARS-CoV-2 challenge. -------------------------------------------------------------27**

**Supplementary Figure S9: ADE of dengue virus infection by the monoclonal and polyclonal antibodies and serum against SARS-CoV-2 in U937 cells------------------28**

**Supplementary Figure S10: Time point-based kinetics of ADE of dengue virus infection by the SARS-CoV-2 anti-Spike monoclonal antibodies and sera raised in mice and hamsters against Spike--------------------------------------------------------------29**

**References:-------------------------------------------------------------------------------------- 30**

**Supplementary Table S1: Details of antibodies for computational study**

| Sr. No. | Antibody | PDB-ID | Resolution (Å) | Neutralization status | kD | target | References |
|---|---|---|---|---|---|---|---|
| 1 | Fab14 antibody | 7WPV | 2·46 | Neutralizing | 4·14nM | RBD | [1] |
| 2 | Fab fragment of FC05 antibody | 7D4G | 3·90 | Neutralizing | 4·322 nM | NTD | [2] |
| 3 | Fab fragment of 8D2 antibody | 7DZX | 3·53 | Non-Neutralizing (Infection enhancing Ab) | NA | NTD | [3] |
| 4 | Fab fragment of 2490 antibody | 7DZY | 3·60 | Non-Neutralizing (Infection enhancing Ab) | NA | NTD | [3] |
| 5 | nCoV617 antibody | 7E3O | 2·51 | Neutralizing | 4·23 nM | RBD | [4] |

| | | | | | | | |
|---|---|---|---|---|---|---|---|
| 6 | P36-5D2 antibody | 7FAF | 3·69 | Broadly neutralizing | 6·91 nM | RBD | [5] |
| 7 | Fab fragment of S2E12 antibody | 7K45 | 3·70 | Neutralizing | 1·6 nM | RBD | [6] |
| 8 | WRAIR-2151 antibody | 7N4M | 3·79 | Neutralizing | <1 pM | RBD | [7] |
| 9 | UT28K antibody | 7X7O | 3·75 | Broadly neutralizing | NA | RBD | [8] |
| 10 | Fab fragment of Ab188 antibody | 7X8Z | 4·10 | Neutralizing | NA | RBD | [9] |
| 11 | Fab fragment of ZCB11 antibody | 7XH8 | 2·99 | Neutralizing | 57·5 pM | RBD | [10] |
| 12 | DMAb 2196 antibody | 8D8R | 4·10 | Neutralizing | NA | RBD | [11] |

| | | | | | | | |
|---|---|---|---|---|---|---|---|
| 13 | Fab fragment (MB.02) of CoV-0213 antibody | 8DZI | 3·50 | Broadly neutralizing | 3·27nM (BA.1) | Omicron RBD | 12 |
| 14 | CR3022 antibody | 6W7Y | 3·30 | Neutralizing | 115nM (Fab) <0·1 nM (IgG) | RBD | 13 |
| 15 | Rabies virus antibody RVA122 | 7U9G | 3·39 | Neutralizing (Rabies virus) | NA | RABV-G | 14 |
| 16 | Fab fragment of C8 antibody | 4UTA | 3·00 | Broadly neutralizing (DENV) | 9 nM | DENV-E | 15 |
| 17 | SARS-CoV-2 RBD in complex | 6W41 | 3·08 | Neutralizing | <0·1nM | RBD | 13 |

| | |
|---|---|
| | with human antibody CR3022 |

**Supplementary Table S2: Details of common epitopes indicated by computational study**

| | Antibodies | | | |
|---|---|---|---|---|
| | DENV-2 | SARS-CoV-2 | | |
| Residues | C8:4UTA | CR3022:6W7Y | S2E12:7K45 | DMAb 2196:8D8R |
| R73 | 1 | 1 | 1 | 1 |
| T70 | 1 | 1 | 0 | 1 |
| S72 | 1 | 1 | 0 | 1 |
| R99 | 1 | 0 | 1 | 1 |
| N103 | 1 | 1 | 1 | 0 |

**Supplementary Table S3: Details of paratope residues involved in interaction of E-protein with CR3022**

| DENV2 E protein | CR3022 | Domain of E protein |
|---|---|---|
| B:Q86 | L:Y31 | Domain II |
| B:S81 | L:Y38 | Domain II |
| B:Q77 | L:K36 | Domain II |
| B:R73 | L:Y38 | Domain II |
| B:S72 | H:S100 | Domain II |
| B:E71 | H:I102 , HS103 | Domain II |
| A:N153 | H:Y27 | Domain I |

**Supplementary Table S4: Clinical data of the 48 patient samples taken for the study.**

| Clinical factors | Total 48 |
|---|---|
| First interval | 21 |
| Second interval | 17 |
| Third interval | 10 |
| **Gender** | |
| Male | 27 (56·25%) |
| Female | 21 (43·75%) |
| **Age (Mean)** | 41 yrs |
| Below 18 yrs | 1 |
| 18-35 yrs | 18 |
| 36-60 yrs | 23 |
| 61 above | 6 |
| **Symptoms** | |
| Symptomatic | 40 |
| Asymptomatic | 8 |
| **Severity** | |
| ICU admission | 6 |
| Oxygen required | 11 |
| Ventilation required | 1 |
| **Vaccination status** | |
| Only 1 dose | 9 |
| Both doses | 35 |
| Non vaccinated | 4 |

**Supplementary Table S5: Neutralizing antibody titers of the commercial SARS-CoV-2 antibodies, animal sera, and convalescent plasma samples against the original Wuhan strain, obtained from live virus micro-neutralization assay.** For the antibodies, sera and plasma samples in the first and second intervals, reciprocal of the highest dilution at which no cytopathic effect is observed is taken as nAb titre. For samples collected in the third interval, $FRNT_{50}$ values are reported.

| **Antibodies and Serum samples** | | |
|---|---|---|
| **ID** | **SNT titer against Wuhan** | **Remarks** |
| 4G2 | 10 | Negative |
| M1B | 10 | Negative |
| P1R | 10 | Negative |
| M4B | 10 | Negative |
| MM41 | 40 | Positive |
| 1A9 | 10 | Negative |
| Mser | 320 | Positive |
| Hser | 320 | Positive |
| Rabishield®-100 | 10 | Negative |
| **First interval** | | |
| **ID** | **SNT titer against Wuhan** | **Remarks** |
| #13 | 1280 | Positive |
| #15 | 640 | Positive |
| #18 | 160 | Positive |
| #22 | 2560 | Positive |

| | | |
|---|---|---|
| #27 | 320 | Positive |
| #141 | 1280 | Positive |
| #143 | 20 | Negative |
| #144 | 160 | Positive |
| #150 | 2560 | Positive |
| #151 | 1280 | Positive |
| #152 | 160 | Positive |
| #153 | 40 | Positive |
| #154 | 640 | Positive |
| #155 | 2560 | Positive |
| #164 | 40 | Positive |
| #170 | 160 | Positive |
| #171 | 320 | Positive |
| #177 | 640 | Positive |
| #188 | 2560 | Positive |
| #189 | 160 | Positive |
| #190 | 2560 | Positive |
| **Second interval** | | |
| **ID** | **SNT titer against Wuhan** | **Remarks** |
| #91 | 320 | Positive |
| #92 | 10 | Negative |
| #93 | 40 | Positive |
| #95 | 10 | Negative |

| | | |
|---|---|---|
| #96 | 160 | Positive |
| #97 | 640 | Positive |
| #99 | 20 | Negative |
| #101 | 80 | Positive |
| #103 | 40 | Positive |
| #105 | 20 | Negative |
| #110 | 10 | Negative |
| #111 | 10 | Negative |
| #113 | 160 | Positive |
| #114 | 80 | Positive |
| #115 | 10 | Negative |
| #116 | 20 | Negative |
| #122 | 80 | Positive |
| **Third interval** | | |
| ID | SNT titer against Wuhan | Remarks |
| #1 | 545·55 | Positive |
| #2 | 482·63 | Positive |
| #3 | 258·87 | Positive |
| #4 | 248·2 | Positive |
| #5 | 66·05 | Positive |
| #6 | 195·08 | Positive |
| #7 | 246·06 | Positive |
| #8 | 453·72 | Positive |

| | | |
|---|---|---|
| #9 | 3752·35 | Positive |
| #10 | 124·44 | Positive |

**Supplementary Table S6: Details of antibodies and sera used for the study**

| Sr. No. | Antibody/Serum | Details | Source |
|---|---|---|---|
| 1 | 1A9 | SARS-CoV / SARS-CoV-2 (COVID-19) spike antibody (S2 subunit), Mouse Mab | Genetex<br>(Cat. No. GTX632604) |
| 2 | MM41 | SARS-CoV-2 Spike Antibody, Mouse MAb | SinoBiological<br>(Cat. No. 40591-MM41) |
| 3 | CR3022 | SARS-CoV-2 Spike Protein (CR3022) Human IgG1 mAb (RBD specific) | Cell Signalling Technology<br>(Cat. No. 37475) |
| 4 | M1B | NR-56488<br>Monoclonal Anti-SARS-Related Coronavirus 2 Spike Glycoprotein, Clone 1-3D7 (produced in vitro) | BEI resources |
| 5 | M4B | NR-55296<br>Monoclonal Anti-SARS-Related Coronavirus 2 Spike Glycoprotein Receptor Binding Domain (RBD), Clone 2TP1C3 (produced in vitro) | BEI resources |

| | | | |
|---|---|---|---|
| 6 | M5B | NR-55295<br>Monoclonal Anti-SARS-Related Coronavirus 2 Spike Glycoprotein Receptor Binding Domain (RBD), Clone 2TP1B11 (produced in vitro) | BEI resources |
| 7 | P1R | NR-52947<br>Polyclonal Anti-SARS-Related Coronavirus 2 Spike Glycoprotein (IgG, Rabbit) | BEI resources |
| 8 | Rabishield®-100 | Rabies Human Monoclonal Antibody (rDNA) (100IU)<br>(By Serum Institute of India Ltd.) | Purchased from Pharmacy |
| 9 | 4G2 | Pan flavivirus anti envelope antibody, Mouse MAb | Inhouse purified from Hb112 hybridoma cells |
| 10 | MSer | Mice serum immunized with SARS-CoV-2 Spike | Inhouse |
| 11 | HSer | Hamster serum immunized with SARS-CoV-2 Spike | Inhouse |

**Supplementary Table S7: Primer and probe sequences used of DENV and GAPDH in qRT PCR**

| Gene | | Sequence |
|---|---|---|
| **GAPDH** | Forward primer | 5'- ATTCCACCCATGGCAAATTC- 3' |
| | Reverse primer | 5'- CGCTCCTGGAAGATGGTGAT- 3' |
| | Probe | 5' -(FAM)- AGCTTCCCGTTCTCAGCCTTCAC -(BHQ1)- 3' |
| **DENV** | Forward primer | 5’- GGTTAGAGGAGACCCCTCCC -3’ |
| | Reverse primer | 5’- GGCGTTCTGTGCCTGGA -3’ |
| | Probe | 5’- JOE-CAGGATCTCTGGTCTCTCCCAGCGT–BHQ1 -3’ |

**Supplementary Table S8: Primer and probe sequences used in qRT PCR for animal tissues**

| Gene | Forward Primer (5'-3') | Reverse Primer (5'-3') |
|---|---|---|
| Mouse GAPDH | ACCACAGTCCATGCCATCAC | TCCACCACCCTGTTGCTGTA |
| DENV-2 | GCAGAAACACAACATGGAACGATAGT | TGATGTAGCTGTCTCCGAATGG |

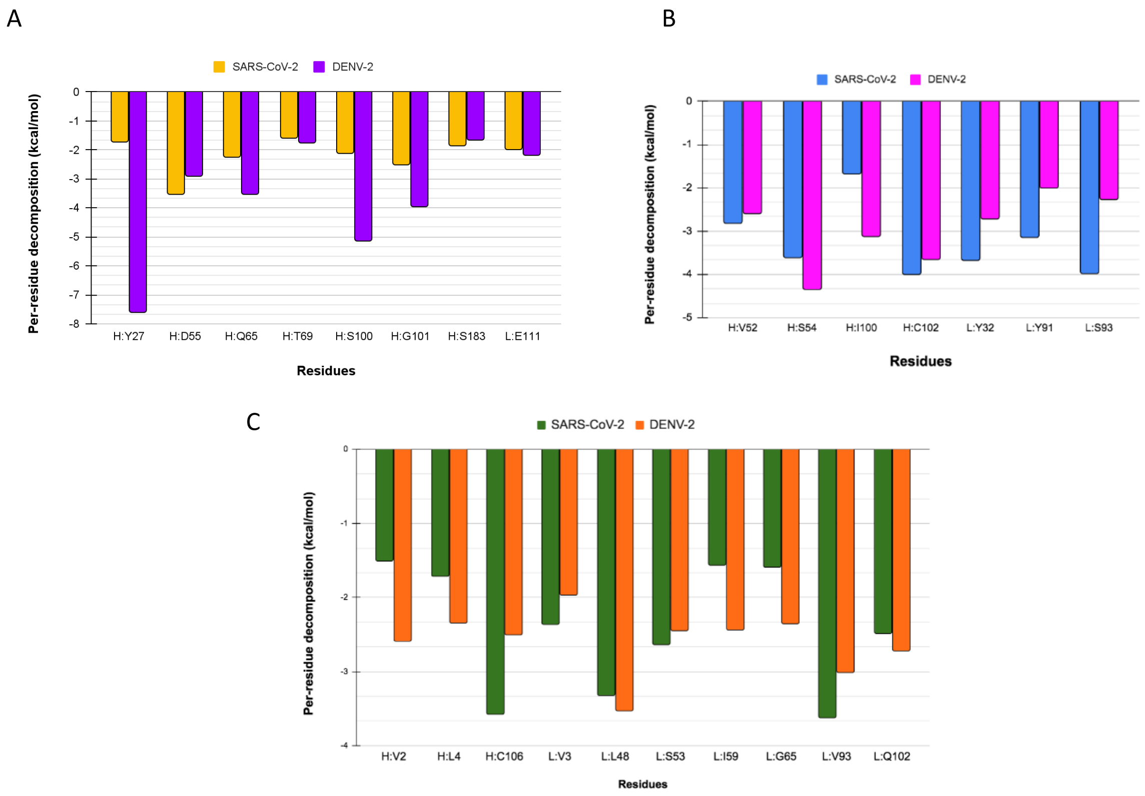

**Supplementary Figure S1: Mining of antibodies interacting residues:** Paratope residues involve interaction with SARS-CoV-2 and DENV-2. (A) CR3022, (B) DMAb 2196, and (C) S2E12. The color code remains the same.

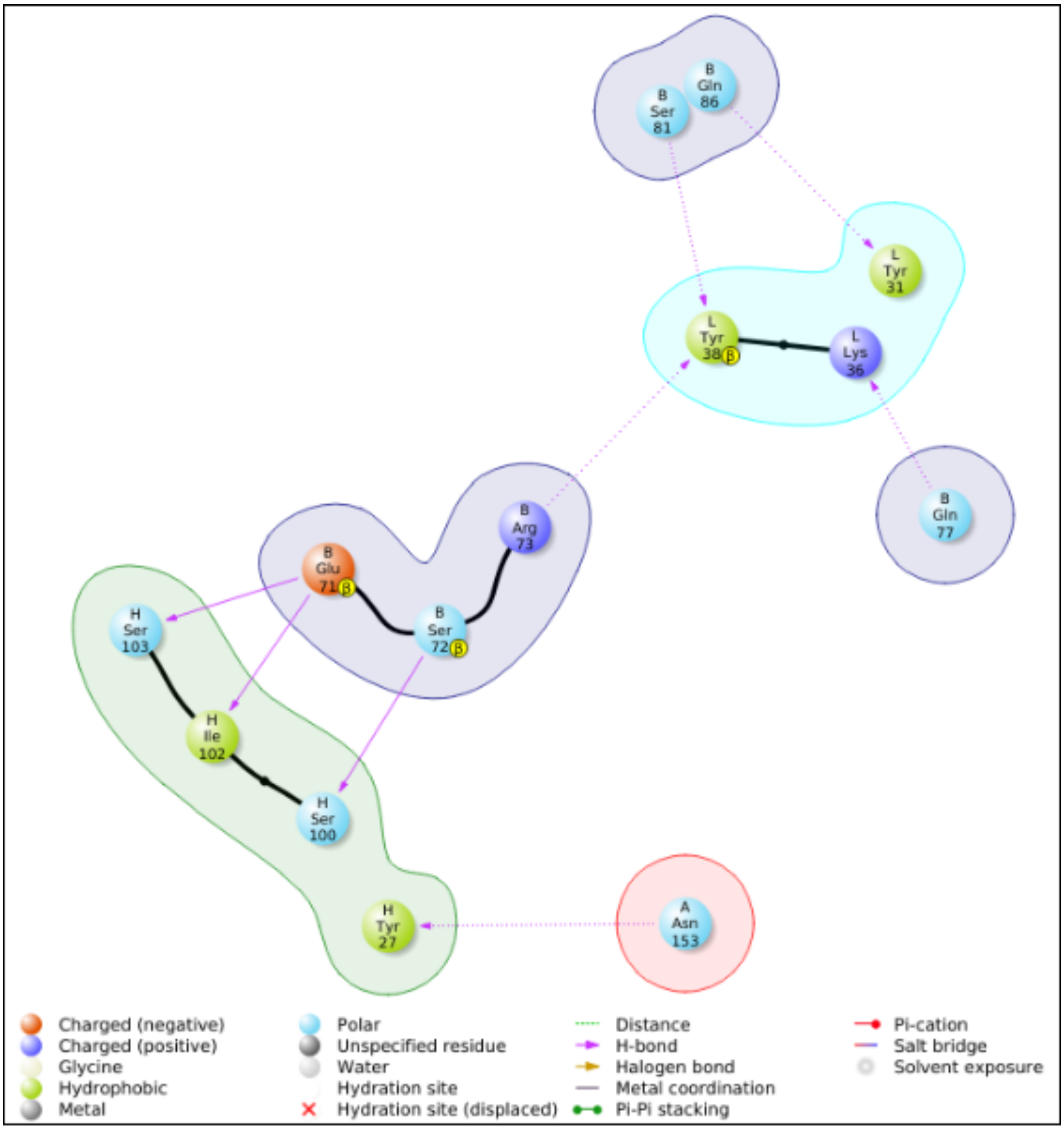

**Supplementary Figure S2: Protein interaction diagram showing paratope residues of mAb CR3022 (chain H L) involved in interaction with E-protein of SARS-CoV-2 (Chain A B).**

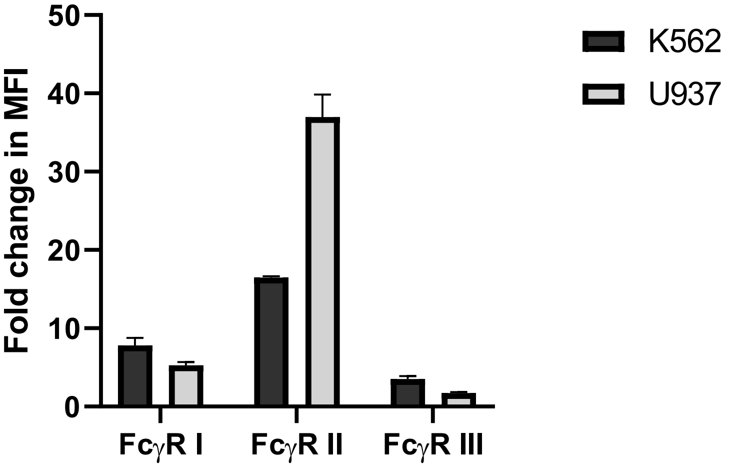

**Supplementary Figure S3: Expression of Fcγ Receptors (FcγRs), type I, II and III in undifferentiated K562 and U937 cells. K562 and** U937 cells were stained with Alexa flour 700 conjugated anti-CD64, PE conjugated anti-CD32, and APC conjugated anti-CD16 antibodies, for assessing the expression of FcγRI, FcγRII, and FcγRIII respectively. The fold change in MFI was calculated with respect to the unstained cells and plotted as the average MFI with standard deviation.

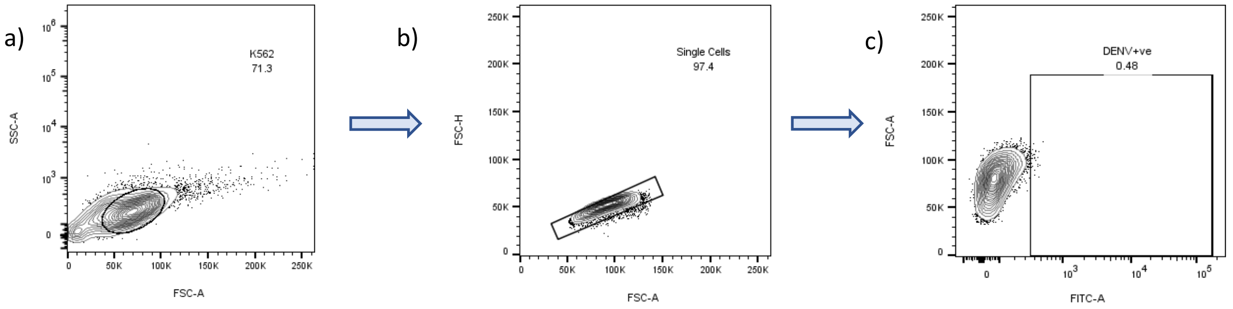

**Supplementary Figure S4: Gating strategy for detection of DENV positive cells in flowcytometry.**

Flowcytometry was used for the detection of Dengue virus-positive cells using pan flavivirus envelope antibody, 4G2 followed by anti-mouse Alexa 488 secondary antibody.

a) Unstained cells were used to set the gates for live population using FSC-A vs SSC-A. b) singlet cells were selected from live population using FSC-A vs FSC-H. c) The singlet cells were used for gating Dengue virus-positive cells using FITC-A vs FSC-A and the percentage of dengue virus positive cells was recorded for further calculations. FITC voltage was set using an isotype control antibody. Data analysis was done on FlowJo v. 10·8·1 and statistical analysis was performed using GraphPad Prism 8·4·2.

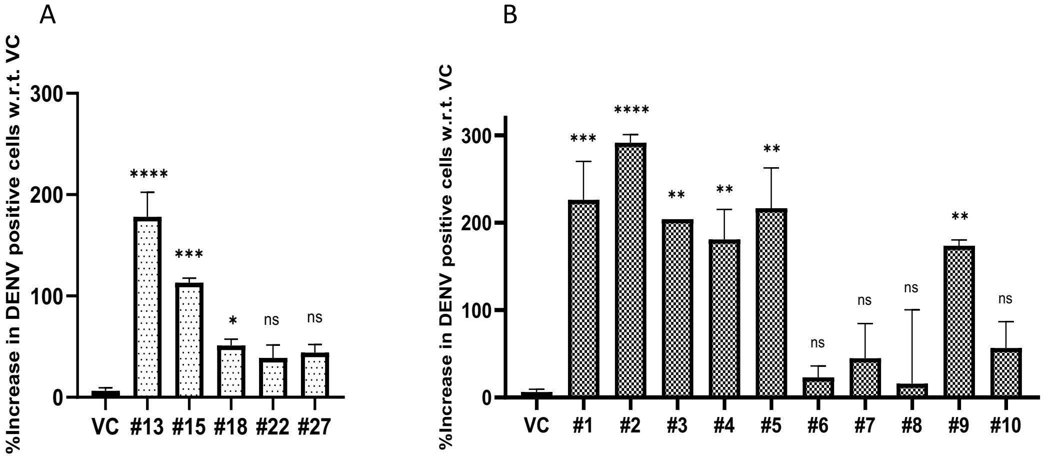

**Supplementary figure S5: ADE of dengue virus infection by the SARS-CoV-2 positive patients' samples in U937 cells.** ADE due to convalescent plasma samples collected in the (A) first interval, n=5 (B) third interval, n=10. Statistical significance was calculated using One-way ANOVA followed by Dunnett's test. Asterisk (*) indicates that the difference between virus control and serum samples is statistically significant. P value = 0·1234 (ns), 0·0332 (*), 0·0021 (**), 0·0002 (***), <0·0001 (****).

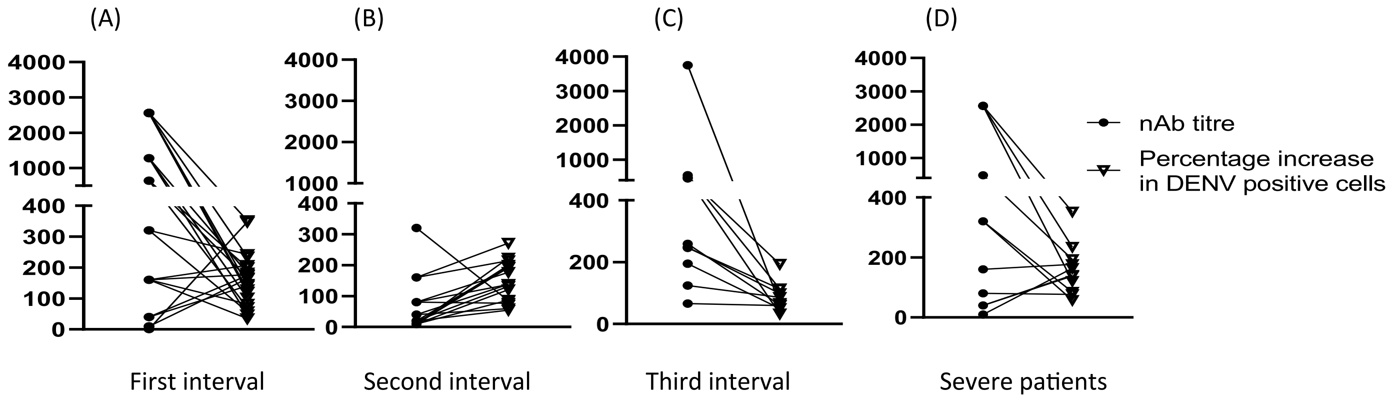

**Supplementary Figure S6: Comparison of neutralizing antibody titres of patient samples against SARS-CoV-2 Wuhan with the increase in dengue virus infection.** nAb titres of patients against SARS-CoV-2 Wuhan were compared to the ability to cause ADE in terms of percentage increase in dengue virus-positive cells in flowcytometry. The comparison was made with samples collected during (A) first interval, n=21 (B)second interval, n=17, (C) third interval, n=10, and (D) Severe patients. The neutralization titres are taken as the reciprocal of the highest dilution showing CPE for samples in first and second intervals. nAb of negative samples (<1: 20) was considered to be 10 Data was analyzed on GraphPad Prism 9·4·1.

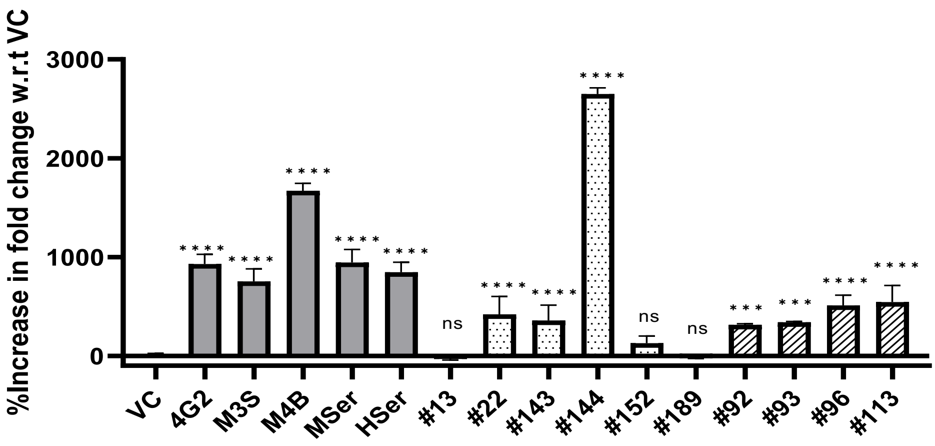

**Supplementary Figure S7: Quantitation of dengue virus ADE using qRT PCR.** ADE assay was performed followed by quantitative real time PCR for detection of DENV2 RNA and the percentage increase in the fold change of DENV-2 RNA copies was measured with respect to the virus control. GAPDH was used as an internal control. Statistical significance was carried out using One-way ANOVA followed by Dunnett's test. Asterisk (*) indicates that difference between the virus control and the serum samples is statistically significant (p < 0·0001). P value = 0·1234 (ns), 0·0332 (*), 0·0021 (**), 0·0002 (***), <0·0001 (****).

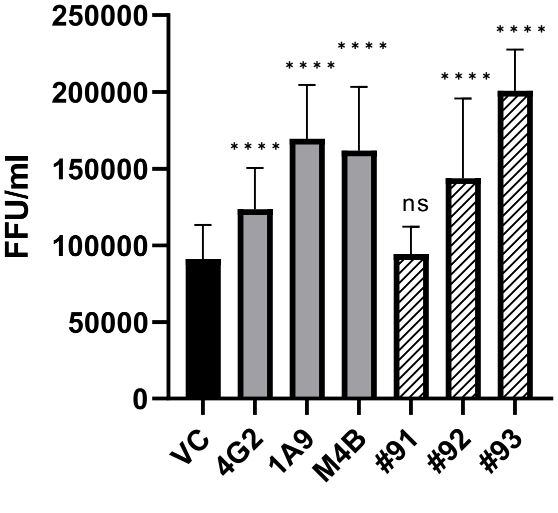

**Supplementary Figure S8: Replicating virus in supernatants from ADE of dengue by the antibodies and convalescent plasma samples against SARS-CoV-2 in K562 cells.** Replicating DENV2 was quantitated as foci forming units/ml (FFU/ml) of the supernatants of ADE assay performed in K562 cells. Serial 10-fold dilutions of the supernatant was used to infect the Vero E6 cells and FFU/ml was calculated 48 h post-infection. Statistical significance was calculated using One-way ANOVA followed by Dunnett's test. Asterisk (*) indicates statistically significant difference between the virus control and treated samples. P value = 0·1234 (ns), 0·0332 (*), 0·0021 (**), 0·0002 (***), <0·0001 (****).

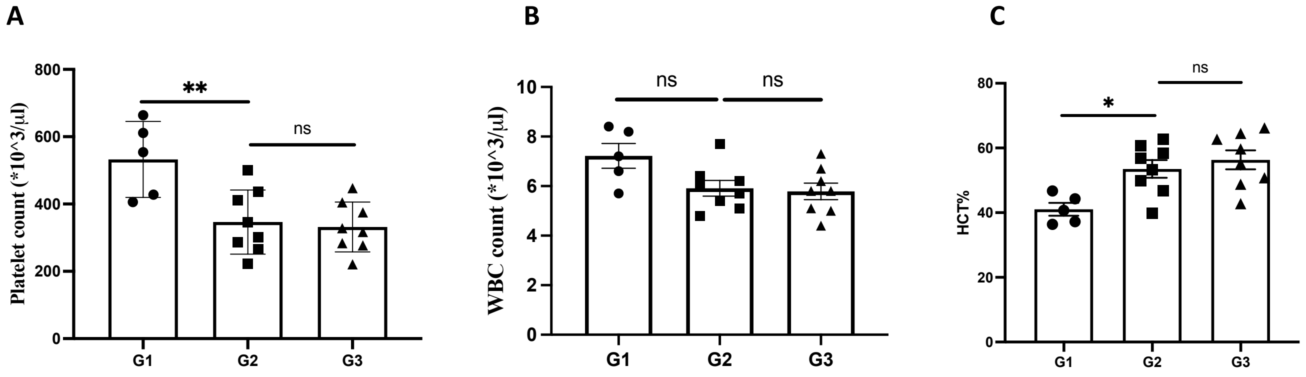

**Supplementary Figure S9. Haematological analysis of dengue-infected AG129 mice following SARS-CoV-2 challenge**. Prefilling blood samples were analyzed for (a) platelet count, (b) WBC count, and (c) haematocrit percentage. Data are presented as mean ± SEM; *P < 0.05, **P < 0.01, ns = not significant.

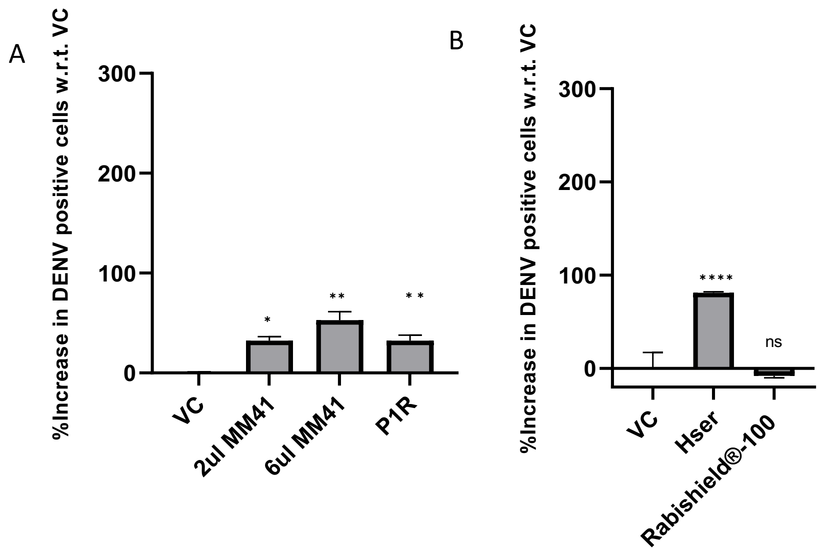

**Supplementary Figure S10: ADE of dengue virus infection by the monoclonal and polyclonal antibodies and serum against SARS-CoV-2 in U937 cells.**

ADE due to (A) 2µl and 6µl of anti-Spike monoclonal antibody MM41 and polyclonal antibody raised in rabbit, P1R, (B) Hamster serum immunized with SARS-CoV-2 spike and negative control (Rabishield®). Statistical significance was carried out using One-way ANOVA followed by Dunnett's test. Asterisk (*) indicates that difference between the virus control and the serum samples is statistically significant ($p < 0{\cdot}0001$). P value = 0·1234 (ns), 0·0332 (*), 0·0021 (**), 0·0002 (***), <0·0001 (****).

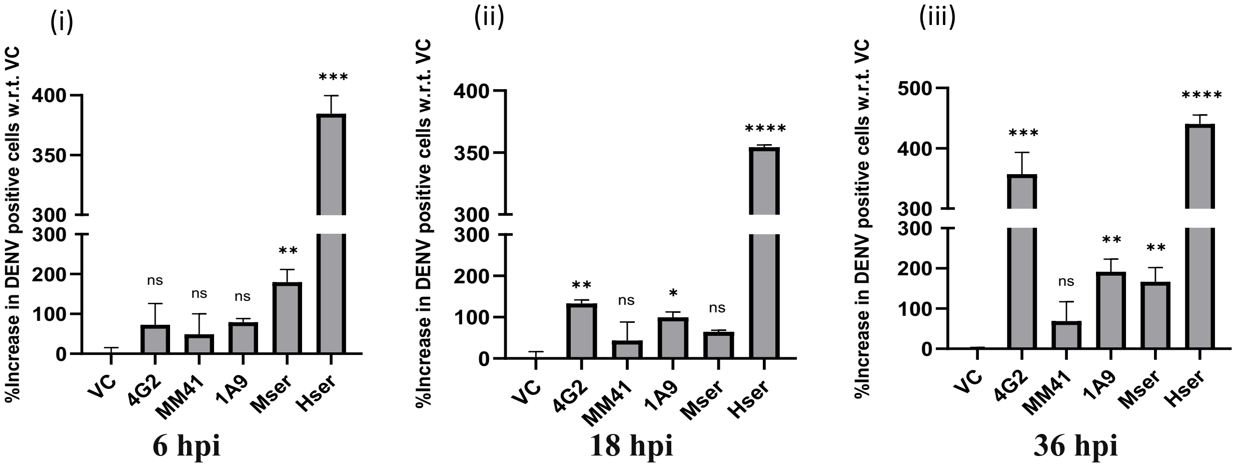

**Supplementary Figure S11: Time point-based kinetics of ADE of dengue virus infection by the SARS-CoV-2 anti-Spike monoclonal antibodies and sera raised in mice and hamsters against Spike.** ADE assay was performed in which post-incubation of antibody and DENV-2, the percentage increase in the DENV-2 positive cells was measured with respect to the virus-only control using flowcytometry. 4G2 was used as a positive Control. (i) ADE due to anti-Spike monoclonal antibodies and animal sera raised against SARS-CoV-2 spike at 6 hpi (ii) 18 hpi, and (iii) 36 hpi. Error bars indicate the standard deviation of each sample performed in duplicates. Statistical significance was carried out using One-way ANOVA followed by Dunnett's test. Asterisk (*) indicates that difference between the virus control and the serum samples is statistically significant (p < 0·0001). P value = 0·1234 (ns), 0·0332 (*), 0·0021 (**), 0·0002 (***), <0·0001 (****).
